# A flexible coding scheme underlying working memory generalization in human parietal and frontal cortices

**DOI:** 10.1101/2025.11.26.690703

**Authors:** Dongping Shi, Luchengchen Shu, Qing Yu

## Abstract

Humans can rapidly extract abstract, common knowledge from distinct experiences, enabling generalization of this knowledge across tasks within working memory (WM) to guide behavior. Although task-specific representations of WM have been observed in a distributed network encompassing sensory, parietal, and frontal cortices, whether these regions construct task-generalizable representations and whether such representations can be dynamically shaped by task demands remain unclear. Across two functional MRI experiments, we investigated WM generalization across distinct WM tasks (location and object tasks) based on shared underlying task structures. Trial-specific task demands varied between passive maintenance and active, rule-guided manipulation of mnemonic stimuli. The posterior parietal cortex (PPC) demonstrated task-generalizable stimulus representations during both maintenance and manipulation. In contrast, the lateral (LFC) and medial (MFC) frontal cortices supported manipulation- and maintenance-based generalization, respectively. Critically, manipulation-based generalization in the PPC and LFC emerged even when participants did not explicitly learn the mapping between task spaces, indicating that active exploration of task structure can spontaneously facilitate generalization across tasks. Together, these findings reveal that flexible generalization for goal-directed behavior is achieved via a distributed WM network, with distinct regions serving active versus passive task demands.

## Introduction

To survive in an ever-changing environment, one must rapidly evaluate the current situation and implement appropriate strategies. This capacity to extract abstract knowledge about the world and apply it to novel situations, known as generalization [1,2], is fundamental to learning and intelligence. Humans excel at this ability, which distinguishes them from other species. Yet, despite growing interest in how generalization is implemented in the brain, many aspects of its underlying mechanisms remain poorly understood.

Successful generalization relies on the abstraction of commonalities across scenarios. While past research has primarily focused on how abstract knowledge or structures are constructed through incremental, long-term learning [3–6], how the brain utilizes these abstraction signals to solve task-specific problems at short, on-task timescales remains less clear. Achieving such rapid computations requires a mental workspace that allows information to be flexibly maintained and manipulated following task rules, commonly known as working memory (WM) [7,8].

Previous studies have demonstrated that task-specific stimulus content in WM is maintained across a distributed cortical network in both humans and non-human primates [9–12]. Particularly, some regions in sensory and lateral frontoparietal cortices can encode multiple stimulus types, forming the basis for generalizable neural codes between different stimuli. Indeed, it has been shown that humans can abstract stimulus information in WM based on shared spatial features [13], and this abstraction is observed neurally in early visual cortex (EVC) and posterior parietal cortex (PPC) [14]. However, generalization can also occur without spatial or visual similarity, and little is known regarding how non-visual generalization is achieved in WM.

Meanwhile, abstract task variables, such as rules that act on stimuli to guide behavior, represent another crucial dimension of WM when stimuli need to be manipulated. Task rule representations primarily engage the lateral frontal cortex (LFC) [15–20]. More recently, it has been found that rule information in WM is also encoded in a distributed cortical network [21]. These task-specific rule signals may interact with task-specific stimulus representations in sensory and lateral frontoparietal regions, resulting in flexible stimulus representations that dynamically adapt to changing task demands during WM [22–24]. These results raised the question of whether task-generalizable representations of stimuli and rules, if exist during WM, are also flexible in dynamic environments. One possibility is that, even when the underlying task structure remains unchanged, specific task demands in WM may shape the level of generalization. For instance, active exploring a task environment versus passive maintaining the same environment would likely require different levels of generalization of task knowledge.

In this study, we address these gaps by systematically examining the neural mechanisms underlying WM generalization. We designed a WM paradigm incorporating two task conditions with flexible task demands: passively maintaining the stimulus versus actively manipulating the stimulus following trial-specific rules [25]. This paradigm was applied to two distinct stimulus sets (location and object), such that both WM tasks shared identical stimulus and rule structures while differing significantly in appearance. Across two experiments, participants performed the location and object tasks either with (Experiment 1) or without (Experiment 2) explicit mapping between task spaces. Leveraging this paradigm, we investigated whether the brain constructs task-generalizable stimulus and rule representations independent of sensory details during WM, how generalization is shaped by task demands, and whether generalization can spontaneously occur without explicit knowledge of the mapping between task spaces. By addressing these questions, we aimed to provide a comprehensive understanding of how the brain implements flexible generalization during WM in a task-dependent manner. We focused our primary analyses on the WM network, but we also examined the medial network such as the medial frontal cortex (MFC), as it has also been implicated in structure learning and cognitive mapping [1,3–5].

## Results

### Behavioral tasks and performance

In both Experiments 1 (N = 23) and 2 (N = 23), human participants performed two WM tasks (a location task and an object task) inside an MRI scanner. In the location task (Figure 1A), stimuli were drawn from a circular location space, and participants mentally maintained or manipulated spatial locations by a cued angle (rotating 0°, ±60°, ±120°, ±180°). In the object task (Figure 1B), stimuli were objects selected from a validated circular object space [26]. Prior to fMRI scanning, participants acquired the structure of this space by learning the transitional relations between objects in the space (Supplementary Figure 1A). During the fMRI session, they mentally maintained or manipulated objects according to symbolic cues indicating the stepwise distances to be updated (0, ±1, ±2, ±3 steps; represented by arrow-shaped cues in Figure 1B). In other words, the two tasks shared similar circular stimulus structures and rule structures but used distinct stimulus sets. Note that the two tasks were performed independently (see Methods). Stimuli were presented in the periphery in the location task, but at the center in the object task. In both tasks, participants maintained central fixation prior to response.

**Figure 1.**
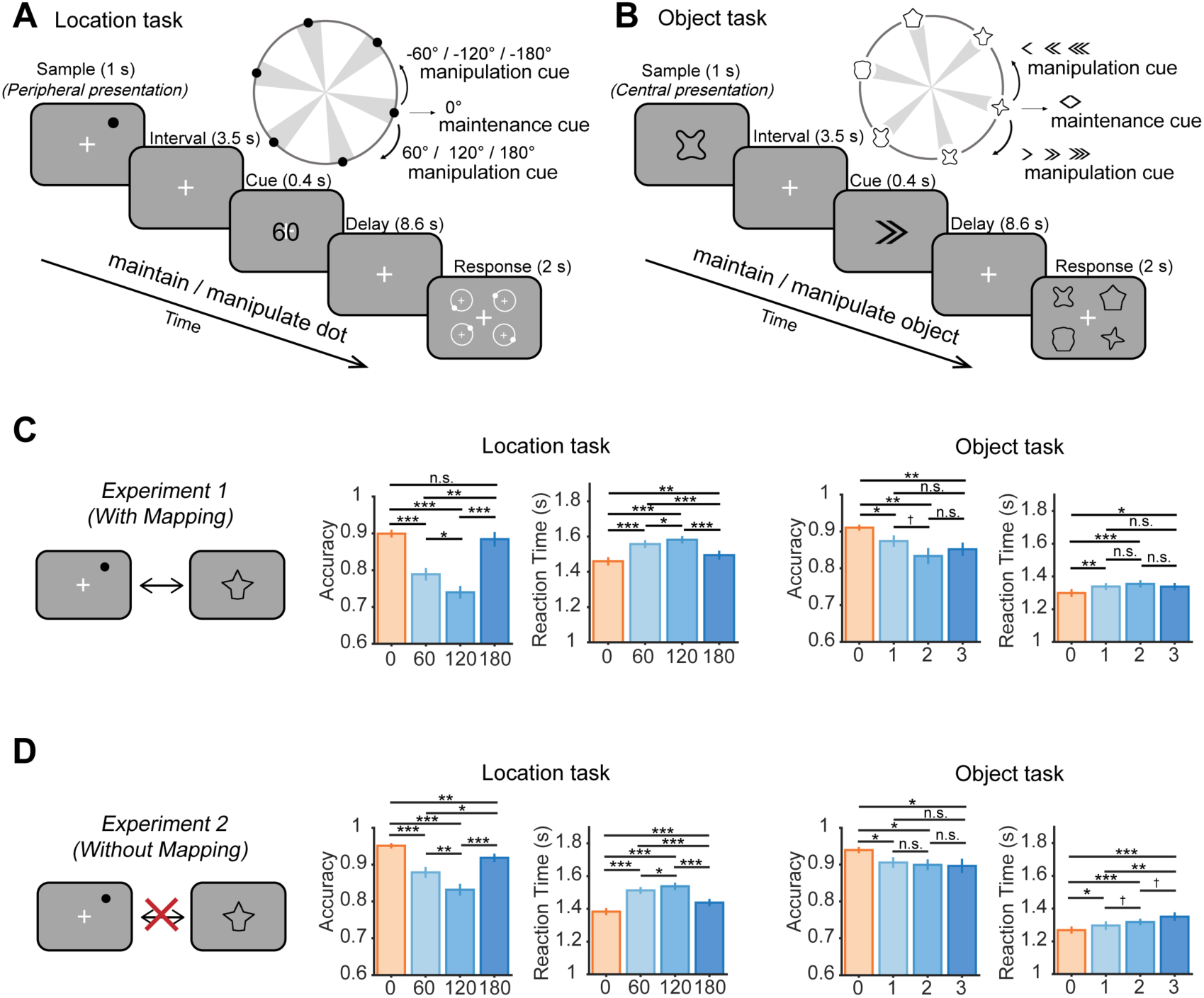
Task paradigm and behavioral performance. (A) Paradigm of the location task. Participants first viewed a dot presented at a peripheral location during the sample period. The dot was randomly chosen from one of 6 bins within a circular location space. A subsequent digit cue indicated whether participants needed to maintain or manipulate the dot location. In the maintenance condition, the cue was always 0, and participants needed to maintain the dot location during the delay period. In the manipulation condition, the cue could be any one of ±60, ±120, or ±180, and participants needed to manipulate (mentally rotate) the dot location either clockwisely (positive values) or counterclockwisely (negative values) by the corresponding degree. During the response period, participants selected the correct dot location from 4 probes. (B) Paradigm of the object task. The object task shares the same task structure as the location task, except that all stimuli and cues were replaced with object versions. During the sample period, an object was presented at the center of the screen, which was randomly selected from one of 6 bins within a validated circular object space. Participants needed to manipulate (mentally transform) the object either clockwisely or counterclockwisely by the corresponding number of steps, and the steps were learned during a behavioral session prior to scanning. During the response period, participants selected the correct object from 4 probes. In both tasks, participants were required to fixate centrally for the duration of each trial before response. (C-D) Behavioral accuracy and reaction time for the location (middle panels) and object tasks (right panels) under different cue magnitudes for Experiment 1 (C) and Experiment 2 (D). Black asterisks indicate FDR-corrected significance, \*\*\**p* < 0.001, \*\**p* < 0.01, \**p* < 0.05, †*p* < 0.1, n.s. *p* ≥ 0.1. Error bars denote ±1 SEM.

Experiments 1 and 2 followed the same task procedure, differing only in whether an explicit mapping between the two stimulus spaces was learned prior to fMRI scanning. Specifically, in Experiment 1, participants were trained to form a fixed one-to-one mapping between location and object stimuli (Supplementary Figure 1B), such that each location was associated with a unique object (Figure 1C); whereas in Experiment 2, this training was absent (Figure 1D).

For both experiments, we analyzed behavioral accuracy and reaction time according to cue magnitude (i.e., rotation angle or step size; Figures 1C-D). Behavioral performance systematically varied as a function of cue magnitude: behavioral accuracy decreased and reaction time increased with greater cue magnitude (excluding 180° in the location task or step-3 in the object task), indicating that participants were actively engaged in the manipulation tasks. To quantify this trend, we fitted linear mixed-effects models (LMEM) and found significant negative relationships between cue magnitude and behavioral accuracy (Experiment 1: location task: *β* = -0.080, *F*(1,67) = 67.878, *p* < 0.001; object task: *β* = -0.038, *F*(1,67) = 19.718, *p* < 0.001; Experiment 2: location task: *β* = -0.060, *F*(1,67) = 67.721, *p* < 0.001; object task: *β* = -0.020, *F*(1,67) = 10.884, *p* = 0.002), as well as significant positive relationships between cue magnitude and reaction time (Experiment 1: location task: *β* = 0.061, *F*(1,67) = 119.35, *p* < 0.001; object task: *β* = 0.028, *F*(1,67) = 28.407, *p* < 0.001; Experiment 2: location task: *β* = 0.077, *F*(1,67) = 128.26, *p* < 0.001; object task: *β* = 0.025, *F*(1,67) = 24.956, *p* < 0.001).

In addition, a closer examination on the behavioral performance revealed an intriguing change with mapping training. The linear relationship between cue magnitude and behavioral performance was violated in trials with the largest cue magnitude (i.e., 180° in the location task or step-3 in the object task). This was likely because participants adopted a different strategy for these trials to facilitate responses, such as mentally flipping the stimulus. However, this flipping strategy was asymmetric between tasks. Specifically, when the two task spaces were not explicitly linked, flipping was observed in the location task but not the object task. In contrast, when the two task spaces were trained to be associated, this pattern became evident in both tasks. LMEM confirmed changes in strategy for object tasks between Experiment 1 and 2, with a significant interaction effect between experiment and reaction time (*β* = 0.014, *F*(1,180) = 5.262, *p* = 0.023). As a control, this interaction effect was not observed in location tasks (*β* = 0.006, *F*(1,180) = 0.452, *p* = 0.502). The interaction effects were absent between experiment and accuracy (object task: *β* = 0.008, *F*(1,180) = 1.186, *p* = 0.278; location task: *β* = -0.005, *F*(1,180) = 0.198, *p* = 0.657). These results suggest that training on explicit mapping likely changed participants’ strategy in performing the object task, such that behavioral strategies between tasks became more assimilated as a result of training, effectively promoting generalization between task spaces.

### IPS as a stimulus-general brain region for working memory

Having observed that mapping facilitated the level of generalization between tasks during WM, we next sought to understand how stimulus information was generalized at the neural level, when the two tasks were explicitly linked in Experiment 1. One critical prerequisite for stimulus generalization is representations of both stimulus domains within the same brain region. For this purpose, we employed representational similarity analysis (RSA) to decode stimulus representation in location and object tasks. The RSA used a continuous model RDM, which assumed that neural dissimilarity increased as a function of stimulus similarity. Within each task, the maintenance condition involved retaining a single stimulus, whereas the manipulation condition resulted in an additional stimulus after mental transformation. For clarity, we refer to the stimulus before transformation as the original stimulus, and the stimulus after transformation as the transformed stimulus in the manipulation condition (Figure 2A). We first focused on three brain regions of interest (ROIs) that have been extensively studied in WM, including EVC, intraparietal sulcus (IPS) in the PPC, and superior precentral sulcus (sPCS) in the LFC [10,11,24,27–30].

**Figure 2.**
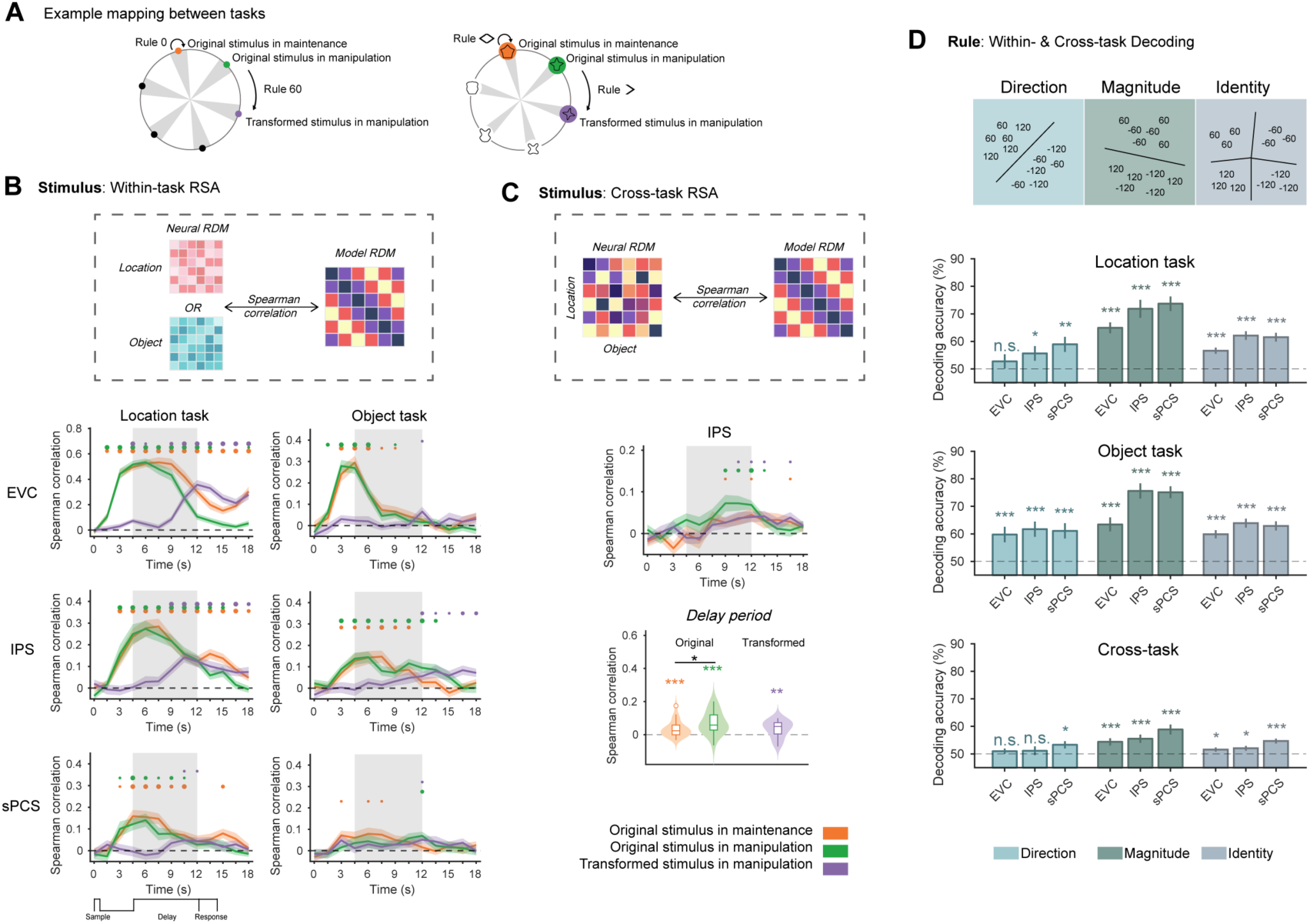
Within-task and cross-task decoding results for stimulus and rule in location and object tasks of Experiment 1 (N=23). (A) Example mapping of stimulus information between location and object tasks in the maintenance and manipulation conditions. (B) Time course of within-task RSA results for stimulus in EVC, IPS, and sPCS, for location (left panels) and object (right panels) tasks separately. The time axis reflects raw time points after trial onset, unadjusted for hemodynamic response delay. The gray shaded areas indicate the delay period. Horizontal dots on top indicate FDR-corrected significance for each condition over time, with dot sizes indicating different levels of significance: large (*p* < 0.001), medium (*p* < 0.01), and small (*p* < 0.05). Error bars denote ±1 SEM. (C) Time-course (middle panel) and epoch-averaged (bottom panels) cross-task RSA results for stimulus in IPS. Violin plots in the bottom panels show the probability density of data (shaded area) with the interquartile range (box), median (inner line), data range excluding outliers (whiskers), and outliers (hollow dots). Epochs were averaged for the delay period (9 – 12 s). (D) Within-task and cross-task SVM decoding results for cue direction, magnitude, and identity in EVC, IPS and sPCS, averaged across the delay period. Error bars denote ±1 SEM. Asterisks on top indicate FDR-corrected significance. \*\*\**p* < 0.001, \*\**p* < 0.01, \**p* < 0.05, n.s. *p* ≥ 0.05.

In the location task, we observed significant and persistent decoding for the original stimulus in both maintenance and manipulation conditions throughout the delay period, in all three ROIs (Figure 2B left panel). Representation of the transformed stimulus also emerged during the middle of the delay period in EVC and IPS, but was substantially weaker in sPCS. Note that the early decoding of the transformed stimulus at the onset of the delay in EVC (Figure 2B left panel) likely resulted from an anticorrelation between the original and transformed stimuli in the manipulation condition (i.e., the original and transformed stimuli cannot be the same), and disappeared when this anticorrelation was controlled for (Supplementary Figure 2). In the object task, robust object representation was still observed in IPS for both original and transformed stimuli (Figure 2B right panel). By contrast, EVC showed significant object representation during memory encoding but decayed quickly during delay, suggesting that EVC was less engaged in object WM. Moreover, object representation in sPCS remained weak (Figure 2B right panel). These results further suggest that participants performed the two tasks independently, rather than simply recoding objects into locations, because if the latter were the case, EVC should have shown robust stimulus decoding towards the end of the object task [31,32]. Combined, these results demonstrate that IPS serves as a stimulus-general brain region for maintaining and manipulating stimulus information during WM.

### Enhanced stimulus generalization through manipulation in IPS

Having established that IPS encoded both location and object WM, we next asked whether information could be generalized across two stimulus domains. This was implemented by performing a cross-task RSA based on learned mapping to evaluate similarities between stimulus-specific neural activity of the two tasks (Figure 2C upper panel). We found significant cross-task decoding for all three stimulus types in the IPS (Figure 2C middle panel). Notably, enhanced cross-task representation was observed for the original stimulus in the manipulation condition compared to maintenance, despite that these two stimuli remained visually identical during sample presentation. Importantly, this generalization pattern emerged during memory delay (*p*s < 0.003 for stimulus decoding, *p* = 0.017 for difference between maintenance and manipulation; Figure 2C lower panel) and was absent during memory encoding, suggesting that it reflected WM processes rather than automatic associative responses evoked by visual presentation. These findings were replicated using support vector machine (SVM) decoders (Supplementary Figure 3), and were consistent across IPS subregions (IPS0-5) (Supplementary Figure 4). Moreover, we confirmed that the difference in cross-decoding between manipulation and maintenance cannot be simply explained by activation differences between conditions [33,34] (Supplemental Figure 5).

### Rule generalization in frontal but not parietal cortex during WM

If stimulus generalization during WM primarily engages the IPS, would the same brain region also be responsible for rule generalization? To address this question, we performed within-task and cross-task SVM decoding for rule information. Multiple dimensions of rule information were present in the current task design: the rule dimension that distinguishes between different cue magnitudes, that distinguishes between different cue directions (clockwise vs. counterclockwise), and that combines the magnitude and direction dimensions (cue identity). We argue that cue direction serves as the optimal choice for rule generalization, as this information is specific to the task structure and cannot be simply explained by alternative factors such as task difficulty; but we nonetheless include rule generalization results using other rule dimensions for completeness (also see Supplementary Figure 6 for the behavioral performance by cue identity and direction). Note that we focused on manipulation-based rules only for all subsequent analyses.

Within each task, cue direction could be reliably decoded in IPS (*p*s < 0.031) and sPCS (*p*s < 0.005), but not in EVC (*p* = 0.146) during the location task. However, in stark contrast with stimulus generalization, cross-task SVM revealed that cue direction was generalizable between tasks in the sPCS (*p* = 0.023), but not in the IPS (*p* = 0.239) or EVC (*p* = 0.239). Moreover, although cue magnitude and identity were decodable in all three regions, cross-decoding performance was also the highest in the sPCS compared to the two other regions (Figure 2D). These results suggest that stimulus and rule generalization in WM are supported by separate neural substrates, with stimulus generalization primarily engaging the IPS and rule generalization primarily involving the sPCS.

### Whole-brain characterization of stimulus and rule generalization

To provide a more comprehensive understanding of stimulus and rule generalization beyond the IPS and sPCS, we performed two additional analyses. First, given the long-established role of the frontal cortex in WM [8,15,35], we parcellated the frontal cortex following previous literature [36,37] into 12 ROIs (Figure 3B), including the premotor cortex (PMC), lateral prefrontal cortex (PFC), and MFC, and performed stimulus and rule decoding in each ROI (Figure 3A). In general, lateral frontal ROIs that exhibited the most robust generalization effects in the manipulation condition were those that lie closely (or overlappingly) with sPCS. Second, to identify potential brain regions exhibiting stimulus or rule generalization at the whole-brain level, we performed whole-brain searchlight analyses. The searchlight results were largely consistent with the ROI analyses, with IPS and LFC subregions (including sPCS and lateral PFC) demonstrating enhanced generalization during memory manipulation and sPCS demonstrating robust cue direction generalization (Figure 3C-D). Intriguingly, in both analyses, we found that in contrary to IPS and sPCS, subregions in the MFC, including medial PFC and anterior cingulate cortex (ACC), showed robust stimulus generalization during WM maintenance but not during WM manipulation (see Supplementary Figure 7 for additional discussion). Similar results were also observed in bilateral middle cingulate cortex (MCC) and right insula in the searchlight analyses. Importantly, the strength of generalization signals in IPS and frontal subregions predicted behavioral RT at the within-subject level, indicating their functional significance in facilitating performance on relevant WM conditions (Supplementary Figure 8). These results together suggest a functional dissociation between the lateral and medial networks during WM generalization: the lateral frontoparietal network primarily supports generalization during active manipulation processes, whereas the medial network contributes predominantly to maintenance-based generalization. Searchlight results for each stimulus type and other cue dimensions can be found in Supplementary Figure 7.

**Figure 3.**
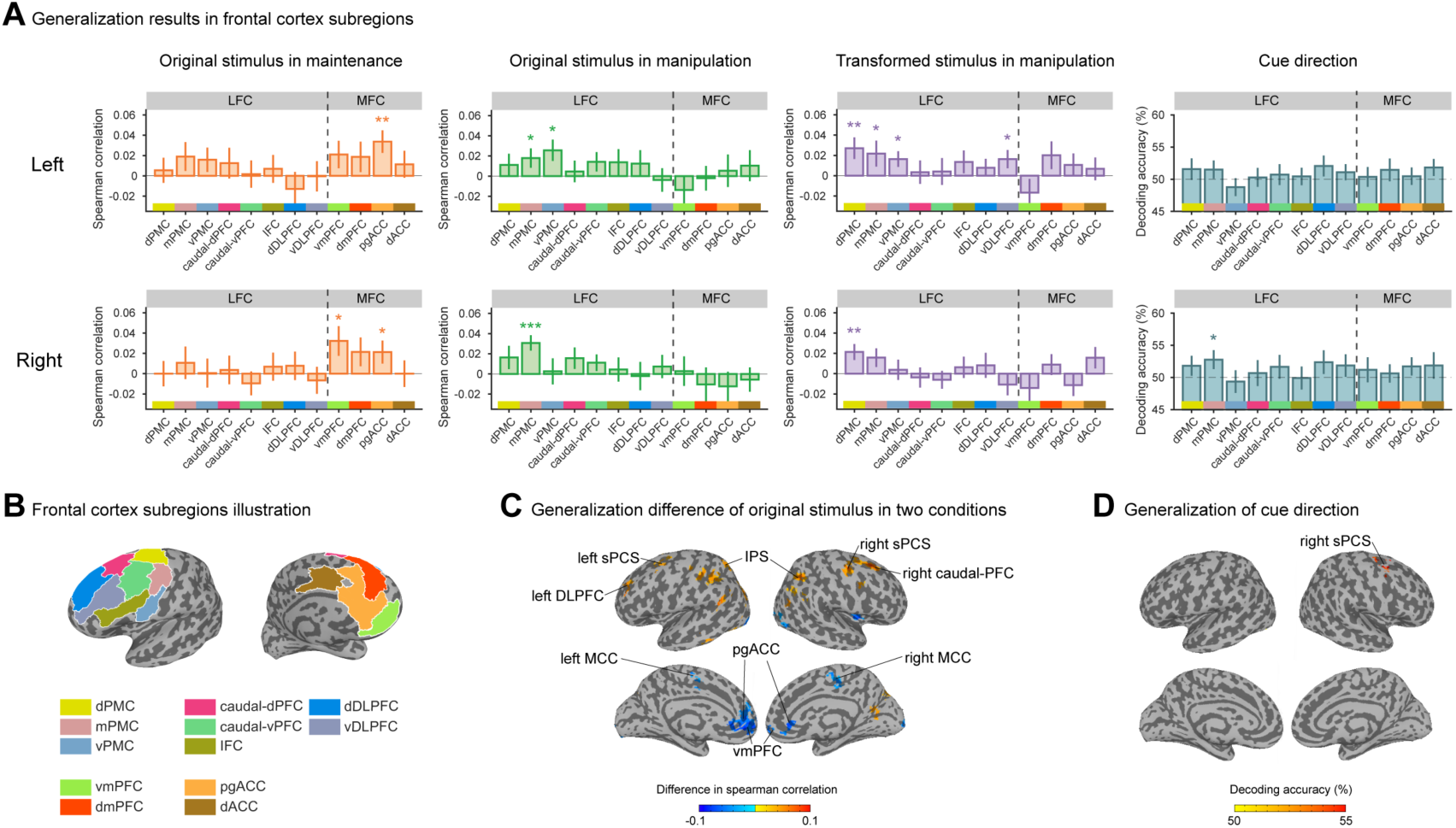
Cross-decoding results between location and object tasks for stimulus and rule in Experiment 1 (N=23). (A) Cross-task decoding results of three stimulus types and cue direction across frontal subregions (top: left hemisphere; bottom: right hemisphere). All results remain uncorrected. Error bars denote ±1 SEM. \*\*\**p* < 0.001, \*\**p* < 0.01, \**p* < 0.05. (B) Illustration of frontal subregions in (A). (C) Whole-brain searchlight RSA results for cross-decoding differences between the original stimuli during maintenance and manipulation. Red denotes greater cross-task RSA results for original stimulus during manipulation compared to maintenance, whereas blue denotes the opposite. (D) Whole-brain searchlight SVM results for cue direction. Cue direction generalization results were thresholded using significant self-decoding results. Cluster sizes are thresholded at 50 voxels for all searchlights for demonstration purposes. LFC = lateral frontal cortex; MFC = medial frontal cortex; dPMC = dorsal premotor cortex; mPMC = middle premotor cortex; vPMC = ventral premotor cortex; caudal-dPFC = caudal dorsal prefrontal cortex; caudal-vPFC = caudal ventral prefrontal cortex; IFC = inferior frontal cortex; dDLPFC = dorsal dorsolateral prefrontal cortex; vDLPFC = ventral dorsolateral prefrontal cortex; vmPFC = ventromedial prefrontal cortex; dmPFC = dorsomedial prefrontal cortex; pgACC = pregenual anterior cingulate cortex; dACC = dorsal anterior cingulate cortex; DLPFC = dorsolateral prefrontal cortex; caudal-PFC = caudal prefrontal cortex; MCC = middle cingulate cortex.

### Increased level of neural generalization with task space mapping

In the behavioral analyses, we demonstrated that training on mapping facilitated generalization between task spaces. If parietal and frontal regions are crucial for generalization, this facilitation should be observed at the neural level as well. We tested this prediction by comparing the level of neural generalization between Experiments 1 and 2.

Because no explicit link was learned between task spaces in Experiment 2, direct cross-decoding analyses between tasks were not feasible. Nevertheless, within-task decoding could still be performed to investigate stimulus and rule coding in IPS and sPCS (Supplementary Figure 9). Overall, stimulus and rule decoding largely replicated results from Experiment 1: for stimulus decoding, the IPS again demonstrated stimulus-general representations during WM delay, while sPCS showed a weaker pattern. For rule decoding, both IPS and sPCS reliably encoded cue directions within each task. These results confirmed robust stimulus and rule coding in these regions regardless of task mapping.

Next, we quantified the level of generalization between experiments using a state space approach. A common approach to assess neural similarity without using cross-decoding is subspace overlap analysis [38], which characterizes the amount of overlap between neural correlation patterns across tasks. For each experiment, we first performed principal component analysis (PCA) on location data to obtain a low-dimensional location subspace by selecting the first five dimensions, and on object data to obtain a low-dimensional object subspace. We then projected location data into the object subspace and calculated the explained variance of the location data in the object subspace normalized by the variance captured by the location subspace (Figure 4A). Likewise, we calculated the explained variance of the object data in the location subspace. The average explained variance of cross-projections was defined as the alignment index between tasks. A higher alignment index would indicate higher similarity in neural correlation patterns, which is, in other words, a higher level of neural generalization.

**Figure 4.**
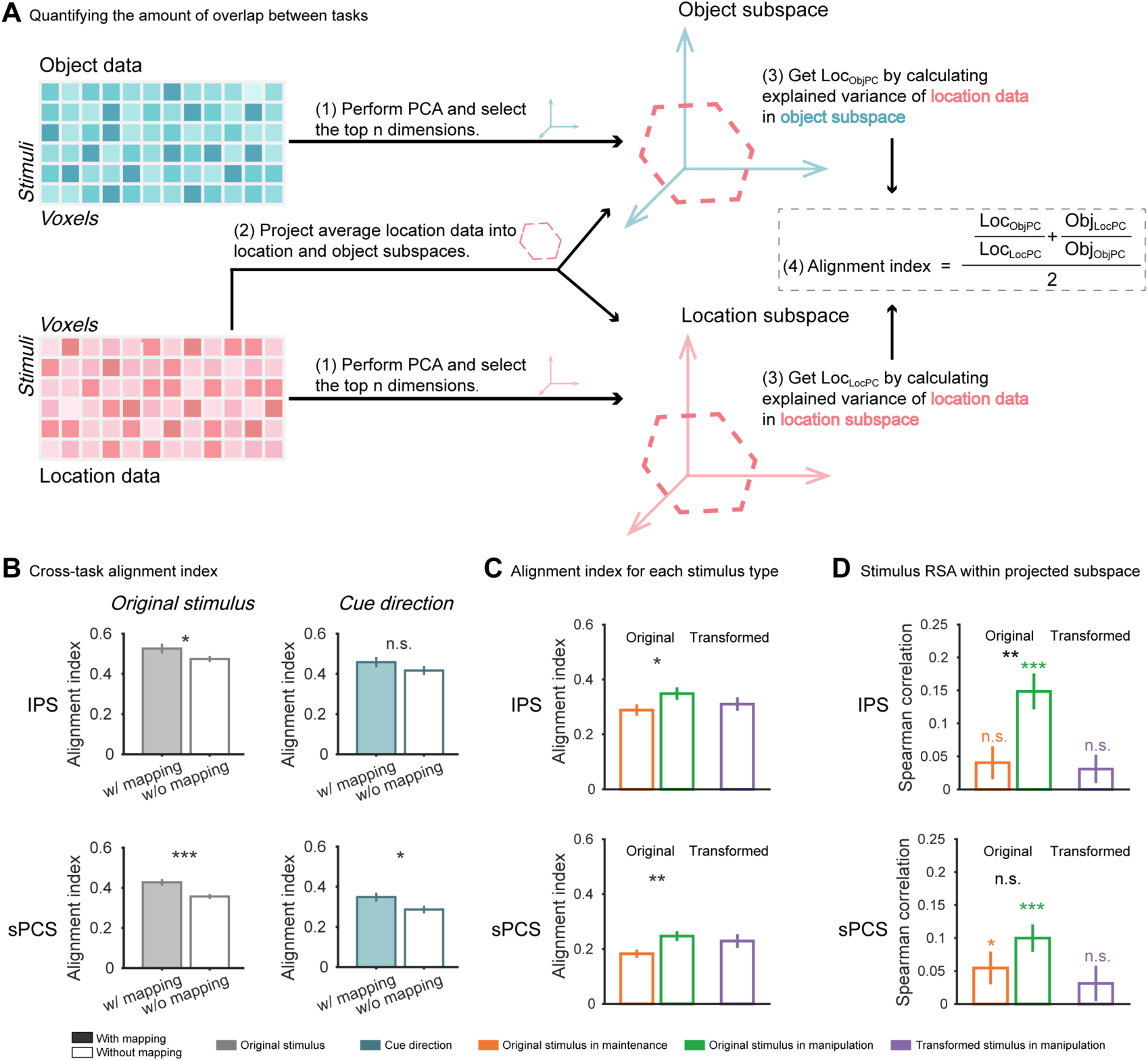
Characterization of generalization using subspace overlap analyses. (A) Schematic for calculating the amount of overlap between tasks (i.e., the alignment index) using the subspace overlap analysis. PCA was performed separately on location and object tasks to obtain location and object subspaces. The explained variance of the location data in the object subspace normalized by the variance captured by the location subspace, and that of the object data in the location subspace, were averaged to obtain the alignment index. (B) Alignment indices between location and object tasks in Experiment 1 (with mapping) and 2 (without mapping) in IPS and sPCS. Left panel shows results combined across the original stimulus conditions, while right panel shows results for cue direction. Error bars denote ±1 SEM. \*\*\**p* < 0.001, \*\**p* < 0.01, \**p* < 0.05, n.s. *p* ≥ 0.05. (C) Alignment indices for each stimulus type in Experiment 2. (D) Within-task stimulus RSA results in the projected subspaces in Experiment 2. All subspace results are based on the delay period (9 – 12 s). w/ mapping: with mapping; w/o mapping: without mapping.

We separately calculated alignment indices for stimulus and rule (see Methods). For stimulus alignment index, we found that the level of generalization increased from no-mapping to mapping condition during memory delay, in both the IPS (*p* = 0.037) and sPCS (*p* < 0.001; Figure 4B left panel). As a comparison, no significant change was observed in the EVC between experiments (*p* = 0.487; Supplementary Figure 10). For rule alignment index, the increase in generalization from no-mapping to mapping was observed in sPCS (*p* = 0.021) but not IPS (*p* = 0.110; Figure 4B right panel) or EVC (*p* = 0.975). Using the first three dimensions for the subspace overlap analysis yielded similar results (Supplementary Figure 11A). Critically, we confirmed that changes in alignment index cannot be explained by differences in BOLD activation (Supplementary Figure 12). These findings supported our hypotheses and further confirmed the roles of IPS and sPCS in generalizing different aspects of information during WM. This generalization was enabled by increased neural correlation patterns as a consequence of mapping.

### Spontaneous WM generalization without explicit mapping between task spaces

In the previous sections, we provided converging evidence for WM generalization as a result of mapping in Experiment 1. However, real-world generalization behaviors do not necessarily require explicit one-to-one mappings between task spaces. As the two tasks shared identical stimulus and rule structures, it is entirely possible that participants perceived this structural similarity and automatically generalized between task spaces without training. To investigate this possibility, following the previous subspace overlap analysis, we found that in Experiment 2, the original stimulus in the manipulation condition showed a higher alignment index compared to the maintenance condition in both IPS (*p* = 0.035) and sPCS (*p* = 0.008; Figure 4C). Additionally, decoding performance was higher in the projected subspace for the original stimulus in the manipulation condition in IPS (*p* = 0.005), but not in sPCS (*p* = 0.115; Figure 4D). These results suggest that neural alignment between tasks was indeed enhanced during active manipulation. Note that in this and all subsequent analyses, we focused on stimulus generalization only, as all comparisons were restricted to conditions within ROIs and no such comparisons were possible for cue direction.

To further elucidate how this neural alignment was implemented, inspired by recent methods [39,40], we decomposed neural data across the two tasks into task-specific and task-general subspaces and examined their relative contributions. Specifically, we decomposed neural activities during the delay period into three orthogonal subspaces: a location-unique subspace, an object-unique subspace, and a shared subspace (Figure 5A). The unique subspaces represented task-specific variance, while the shared subspace captured common variance of both tasks (i.e., the generalization subspace). Therefore, in the absence of explicit mapping in Experiment 2, generalization (if existed) should primarily impact neural activity in the shared subspace that represented both tasks.

**Figure 5.**
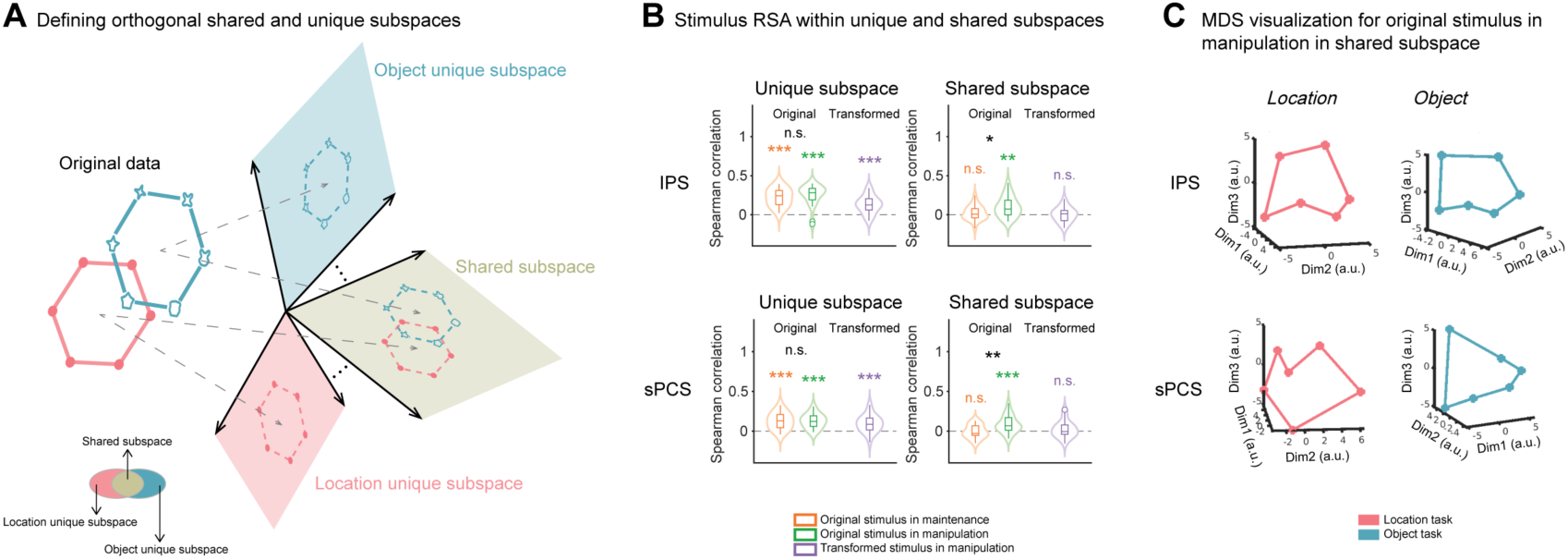
Characterization of generalization using subspace separation analyses. (A) Schematic for extracting orthogonal shared and unique subspaces using the subspace separation analysis. The original data were decomposed into three orthogonal subspaces: location-unique, object-unique, and shared subspaces. The shared subspace represents common variance of both tasks, whereas the unique subspaces only captured individual task variances. The relationships between the three subspaces are also illustrated in the Venn diagram. (B) Within-task stimulus RSA results (average across location and object tasks) in the unique and share subspaces in IPS and sPCS. Violin plots show the probability density of data (hollow area) with the interquartile range (box), median (inner line), data range excluding outliers (whiskers), and outliers (hollow dots). Error bars denote ±1 SEM. \*\*\**p* < 0.001, \*\**p* < 0.01, \**p* < 0.05, n.s. *p* ≥ 0.05. (C) MDS visualization for the original stimulus in the manipulation condition in the shared subspace in IPS and sPCS. All subspace results are based on the delay period (9 – 12 s).

In the IPS shared subspace, we found significant neural representation of the original stimulus in the manipulation condition (*p* = 0.002) but not in the maintenance condition (*p* = 0.203) or for the transformed stimulus in the manipulation condition (*p* = 0.484). Moreover, decoding performance of the original stimulus was significantly higher in the manipulation condition (*p* = 0.029; Figure 5B). In contrast, all stimuli were significantly represented in their respective unique subspaces (*p*s < 0.001), with no difference between conditions (*p* = 0.199). Similar results were also observed in sPCS (Figure 5B), and using SVM decoding (Supplementary Figure 13). These results again suggest enhanced generalization for the original stimulus during manipulation compared to maintenance. Visualization of neural geometry in the shared subspace using multidimensional scaling (MDS) confirmed a circular structure of the original stimulus in the manipulation condition (Figure 5C; MDS results for other stimulus types are shown in Supplementary Figure 14). Additionally, although direct one-to-one mapping between location and object stimuli was not available in Experiment 2, we conducted an exploratory analysis to identify the most likely mapping between tasks by iterating over all possible mappings, and performed cross-task decoding analysis using this mapping. This analysis again revealed higher and more persistent cross-decoding for the original stimulus in the manipulation condition (Supplementary Figure 15). Altogether, these results provide further evidence that active manipulation enhances generalization in the IPS (with a weaker pattern in sPCS), even when the two task spaces were not explicitly linked.

Finally, as a separate check, we compared neural alignment indices between experiments for each stimulus type. This revealed that the increase in generalization was most pronounced in the maintenance condition (IPS: *p* = 0.004; sPCS: *p* = 0.003) rather than manipulation (*p*s > 0.069; Figure 6), suggesting that mapping primarily changed generalization patterns during WM maintenance. This result echoed the above subspace results from Experiment 2, indicating that manipulation-based generalization in IPS and sPCS might reflect a more automatic process that emerged without mapping training, whereas maintenance-based generalization in these regions was more dependent on explicit mapping.

**Figure 6.**
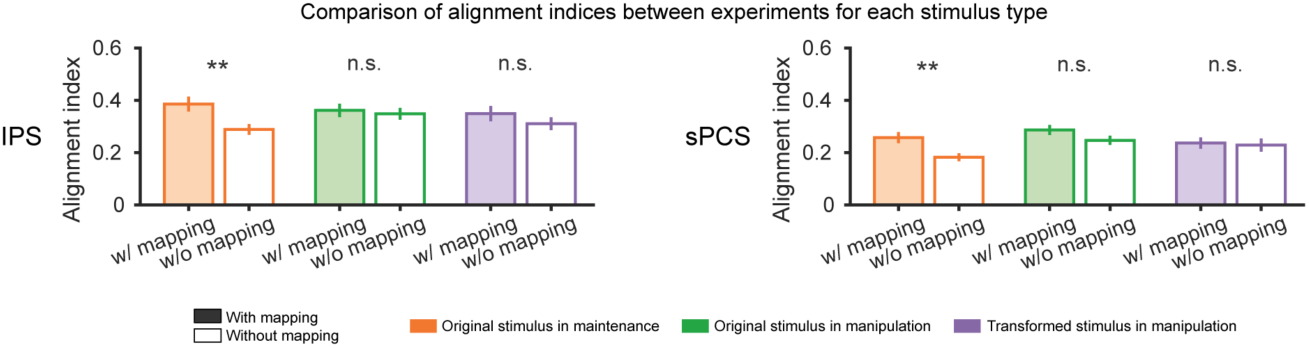
Comparing alignment indices between experiments for each stimulus type. Alignment indices between location and object tasks in Experiment 1 (with mapping) and 2 (without mapping) were plotted for each stimulus type, in IPS and sPCS. Error bars denote ±1 SEM. \*\*\**p* < 0.001, \*\**p* < 0.01, \**p* < 0.05, n.s. *p* ≥ 0.05. All subspace results are based on the delay period (9 – 12 s). w/ mapping: with mapping; w/o mapping: without mapping.

## Discussion

In this study, we investigated how stimulus and rule information are generalized between two WM tasks with shared task structure but distinct stimulus sets. Through two fMRI experiments, we provide converging evidence for three principles of WM generalization. First, the primary neural loci for stimulus and rule generalization in WM are largely spatially separable: stimulus generalization was most robust in the IPS region within PPC, a brain region implicated in WM [41] and structure learning [42]. In contrast, rule generalization primarily engaged the LFC, a brain region critical for WM [8] and cognitive control [15]. These results indicate that WM generalization is highly dependent on the specialized neural modules that actively process the relevant information. Second, WM generalization is highly task-dependent. Manipulation functions of WM, which require active control and online transformation of the maintained stimulus information, facilitated neural generalization in PPC and LFC. Crucially, this enhanced generalization emerged even in the absence of explicit mapping, highlighting its spontaneous nature during active task exploration. Lastly, the neural systems for active versus passive generalization are dissociable. While the lateral frontal network supported generalization during active manipulation, the medial frontal network supported generalization during passive maintenance of task knowledge. Taken together, these findings reveal that rapid, flexible generalization is achieved via a distributed WM network, with distinct cortical regions specialized in different aspects of WM generalization in a task-dependent manner.

The PPC has well-established roles in diverse cognitive functions. Extant evidence from the WM literature suggests that stimulus information held in WM is encoded by delay-period activity in the PPC [11,27,28,41,43]. Compared to the visual cortex, WM representation in the PPC is more task-dependent [28] and is particularly involved in internally-generated processes, such as mental imagery [44] and mental transformation [45,46]. In parallel, the PPC is also implicated in episodic memory-related processes, including memory retrieval [47,48], structure learning [42,49], and cognitive mapping [50,51]. This multifaceted functionality makes PPC an ideal candidate for interactions between WM and episodic memory systems. Such interactions are particularly crucial for WM generalization, as it requires both the retrieval of prior knowledge from long-term memory and on-task implementation of this knowledge. Consistent with this idea, we found that stimulus generalization was most prominent in the PPC. Notably, the level of generalization was highly task-dependent: manipulation demands significantly enhanced stimulus generalization compared to maintenance. This is likely because manipulation facilitated the exploration of the structural relationships between stimuli, leading to better extraction of task regularities and ultimately more efficient task performance through generalized representations. Critically, our task design required trial-by-trial switches between maintenance and manipulation functions, meaning that this difference in generalization emerged rapidly within seconds, rather than slowly developed as different default strategies for different conditions.

Although the work reported here aimed to identify brain regions that can generalize signals irrespective of sensory details, in realistic experimental settings we are often limited to using only a few stimulus domains to test this question. This raises the question of how generalizable our conclusions are to other stimulus domains, task demands, and WM task types [52]. Previous studies have demonstrated that the PPC shows generalizable signals during WM across visual stimuli sharing the same spatial features, either at the same [14,44] or different [53] retinotopic positions. PPC also generalizes WM across stimuli from different sensory modalities sharing the same spatial layouts [22]. Our results extend previous findings by demonstrating that PPC generalizes across stimulus domains even without visual similarity. Importantly, this generalization process differs from associative recoding [31,32], as the lack of object decoding during memory delay in EVC indicates that participants did not simply recode objects into locations to perform the task. Moreover, because stimulus decoding time course in EVC diverged between tasks, generalization was also unlikely to be driven by eye movements, which should have produced synchronized decoding patterns in EVC if present. These findings together suggest that PPC signals generalize across a wide range of stimulus domains and modalities.

In contrast to stimulus generalization, rule generalization primarily involved the LFC, particularly sPCS and its surrounding regions. This finding aligns with the long-established role of the LFC in cognitive control and task-relevant processing [15,19,20,54]. The dissociation between stimulus and rule generalization suggests that WM generalization does not rely on a single brain region, but rather recruits a distributed brain network, with distinct regions responsible for different subfunctions in WM. It is noteworthy that although rule generalization primarily engaged the sPCS in our study, we acknowledge that more anterior prefrontal regions may also contribute to this process, particularly when more complex rules (e.g., hierarchical or schematic control) are recruited [19,20,55].

Previous work has identified a distributed cortical network for representing stimulus and rule information in WM. Here, we demonstrate that within the distributed network, specific brain regions (including PPC and LFC) are critical for stimulus and rule generalization. How should we interpret these generalization signals, which are constrained to specific brain regions, given the presence of widely-distributed WM representations [9–12]? We argue that the current findings help to reconcile the heterogenous nature of these WM representations. While task-specific WM representations enable the flexible execution of individual tasks, generalizable representations reflect higher-order, abstract signals that are related to task knowledge. This division of labor aligns with recent theoretical debates on the dimensionality of neural representations, where high-dimensional representations support flexibility and low-dimensional representations afford stability and generalization [56–58]. How these task-general and task-specific representations interact to facilitate behavior remains a promising future direction to explore.

Why does the brain generate generalization signals in the first place? When explicit mapping was not available between tasks spaces, stimulus generalization was enhanced in the manipulation condition in the PPC compared to the maintenance condition, suggesting that it was by nature a more automatic process when stimulus needed to be manipulated, potentially because task structures were better explored in this context as we discussed earlier. This idea was further supported by the mapping experiment, because when the two task spaces were explicitly linked via training, an increase in the level of generalization was primarily observed during maintenance rather than manipulation. This result suggests that maintenance-based generalization may recruit a more intentional, effortful process, as simple maintenance does not guarantee a thorough exploration of task structure, unless explicitly instructed. These results raise an intriguing possibility that generalization can occur both explicitly and implicitly, the mechanisms of which require further investigation in future research.

The cognitive map literature highlights a primary engagement of the medial temporal lobe and the medial/orbito-frontal (MTL-MOPFC) network in representing physical and abstract task structures [1,3,5,59,60]. These structural representations serve as the foundation for extracting structural similarities and generalization. How should we interpret the current findings in the lateral frontoparietal network as compared to prior work? We propose that the two lines of research can be reconciled when considering their respective roles in task implementation and task knowledge representation [54,61]. Specifically, we observed robust stimulus generalization in subregions of the MTL-MOPFC network (e.g., medial PFC) only during maintenance but not manipulation, suggesting that generalization signals in this network emerged when participants passively maintained the task structure without actively exploring it. Conversely, LFC contributed predominantly to WM manipulation rather than maintenance, indicating its role in active task implementation on stored information. The PPC, as a hub between the two networks, exhibited generalization signals during both maintenance and manipulation. These results together indicate that generalization involves a distributed network beyond the traditional WM or episodic memory systems, with distinct regions serving complementary roles in this process.

In summary, across two experiments, we demonstrate that a distributed cortical network, including the lateral frontoparietal and medial networks, works in tandem to support WM generalization in a flexible, rapid, and task-dependent manner. Whether the current framework extends to other cognitive processes remains to be investigated in future research.

## Methods

### Participants

A total of 47 participants were recruited from the community of Shanghai Institutes for Biological Sciences, Chinese Academy of Sciences. Twenty-four participants (15 females, all right-handed, mean age = 23.9 ± 2.8 years) were recruited for Experiment 1. Among them, 23 participants completed both the location and object tasks, whereas one participant dropped out of the study after completing one task. Twenty-three participants (19 females, all right-handed, mean age = 23.2 ± 1.3 years) were recruited for Experiment 2, all completed both tasks.

All participants had normal or corrected-to-normal vision and normal color vision. They reported being neurologically healthy and were eligible for MRI. All participants provided written informed consent, approved by the ethical committee of Center for Excellence in Brain Science and Intelligence Technology, Chinese Academy of Sciences (CEBSIT-2020028), and were monetarily compensated for their time. The study was conducted according to the principles expressed in the Declaration of Helsinki.

### Stimuli and task procedure

Experimental stimuli were generated and controlled by MATLAB (The Mathworks) and Psychtoolbox-3 extensions [62,63]. The background of the screen remained grey (RGB = 128, 128, 128) throughout the experiment. During fMRI scanning, visual stimuli were presented on a screen with a 60-Hz refresh rate (2560 × 1440 for Experiment 1; 1280 × 1024 for Experiment 2). Viewing distance was 161 cm for Experiment 1, and was 90.5 cm for Experiment 2. Behavioral responses were recorded through a Sinorad MRI-compatible button box.

### Behavioral training

Participants completed the study in three separate sessions on different days. The first session was a behavioral training session where participants acquired the structure of a validated circular object space [26] by learning the transitional relations between objects drawn from the space (Supplementary Figure 1A). After the structure learning task, participants learned the WM tasks to be performed in the following two fMRI sessions. We designed different training protocols for Experiments 1 and 2. In Experiment 1, participants completed behavioral training for both tasks prior to the fMRI sessions. During this training, participants were instructed to form fixed one-to-one mapping relationships between location and object stimuli (Figure 1 & Supplementary Figure 1B). In Experiment 2, all participants performed the location and object tasks in separate fMRI sessions, with task-specific behavioral training conducted prior to the corresponding session. No training on stimulus mapping was introduced in Experiment 2. Behavioral training ended when participants achieved an average accuracy above 85% for each task. The location and object task paradigms are described below.

### Location task

In the location task [25], stimuli were black dots (approximately 0.5° of visual angle) presented at a visual angle of 3.7° from the center of the screen, and were selected from 6 bins (16°, 76°, 136°, 196°, 256° and 316°) with a random jitter of ± 10° in each bin. The task consisted of 2 conditions: maintenance and manipulation. Each trial began with 1-s presentation of a black dot, randomly selected from one of the six bins. Following a 3.5-s interval, a digit cue (1.3° of visual angle) appeared at the center of the screen for 0.4 s. The numerical value of the cue specified the angle to be rotated for the sample dot. Specifically, in the maintenance condition, the digit was always 0, meaning that no mental rotation needed to be performed. In the manipulation condition, the digit was randomly selected from ±60, ±120, or ±180 (with positive values indicating clockwise rotation and negative values indicating counterclockwise rotation). A delay period of 8.6 s followed the disappearance of the cue, during which participants were required to maintain the sample location in the maintenance condition or mentally rotate the sample location by the cued angle in the manipulation condition. At the end of delay, a response screen appeared, consisting of four probes arranged in a 2×2 matrix at the center of the screen, as shown in Figure 1A. Each probe consisted of a circle (2.5° of visual angle) with a dot (0.37° of visual angle) on it. One of the probes contained the correct location of the maintained or rotated location, and dot locations in the three other probes were ±60°, ±120°, or 180° from the correct location. Participants needed to choose the correct probe within 2 s via button presses. Inter-trial intervals (ITI) were equally selected from 4 s, 5.5 s and 7 s and randomized in each run.

Each run consisted of 15 trials, with either 7 maintenance and 8 manipulation trials or 8 maintenance and 7 manipulation trials. The two combinations alternated across runs. Forty-five out of 46 participants completed 16 runs, whereas one participant in Experiment 2 completed 15 runs due to malfunction of MRI scanner. Trials for each location bin were balanced across the entire location task (20 trials for each location in each condition), with the exception of the participant who completed 15 runs. Since the rotation of +180° and −180° resulted in the same location, the trial numbers of these two cues were half of other rotation cues (i.e., 120 trials for 0°, 24 trials each for ±60° and ±120°, 12 trials each for ±180°).

### Object task

The object task shared the same task procedure as the location task, except that all stimuli and cues were replaced with object versions. In the object task, stimuli (3.7° of visual angle) were presented at the center of the screen, and were selected from 6 bins of the learned circular object space, with three objects in each bin. Each trial began with 1-s presentation of an object at the center of screen, randomly selected from one of the six bins. After a 3.5-s interval, a symbolic cue (1.3° of visual angle, see Figure 1B) appeared at the center of the screen for 0.4 s. The cues for maintenance and manipulation were indicated by arrows, with rightward arrows indicating one manipulation direction, leftward arrows indicating the opposite manipulation direction, the number of arrows indicating the manipulation steps, and a diamond-shaped cue (one leftward arrow + one rightward arrow) indicating the maintenance condition. A delay period of 8.6 s followed the disappearance of the cue, during which participants were required to maintain the object in the maintenance condition, or mentally transform the object according to the cue in the manipulation condition. At the end of delay, a response screen appeared, consisting of four objects (each with a visual angle of 2.5°) arranged in a 2×2 matrix at the center of the screen, as shown in Figure 1B. Participants needed to choose the correct probe within 2 s via button presses. ITIs were equally selected from 4 s, 5.5 s and 7 s and randomized in each run.

Trial balancing principles followed those used in the location task. All participants (N = 46) completed 16 runs. In both tasks, a fixation cross (approximately 0.6° of visual angle) was presented at the center of the screen throughout the experiment. Fixation cross was white before response, turned green after participants’ response, and turned black during ITIs.

### fMRI session design

For 8 participants in Experiment 1 (separate-session group) and all participants in Experiment 2, they completed the location and object tasks in separate sessions. Specifically, four from Experiment 1 and twelve from Experiment 2 completed the location task in the first session and the object task in the second session, while the rest completed the object task in the first session, followed by the location task in the second session.

For the rest 15 participants in Experiment 1 (mixed-session group), they performed both the location and object tasks in each session, with task order counterbalanced. Specifically, 8 participants completed the location task in runs 1–8 and the object task in runs 9–16 during the first session, and then completed the object task in runs 1–8 and the location task in runs 9–16 during the second session. The remaining 7 participants followed the opposite sequence.

Participants in Experiment 2 additionally performed an attention task for location and object stimuli, results from this task were not analyzed in the current study.

### Data acquisition

MRI data were collected using a Siemens 3 Tesla MRI scanner (Prisma system for Experiment 1, and Tim Trio system for Experiment 2) with a 32-channel head coil at the MRI Platform of the Brain Imaging Center at Institute of Neuroscience, Center for Excellence in Brain Science and Intelligence Technology, Chinese Academy of Sciences (CEBSIT, CAS). High-resolution anatomical T1-weighted images were acquired using an MP-RAGE sequence (TR = 2300 ms, TE = 2.98 ms, Flip Angle = 9°, 192 slices, Resolution = 1 × 1 × 1 mm). Whole brain functional T2*-weighted images were acquired using a multiband accelerated echo-planar imaging 2D (MBEP2D) sequence (Multiband Factor = 2, TR = 1500 ms, TE = 30 ms, Flip Angle = 60°, Matrix Size = 74 × 74, 46 slices, Resolution = 3 × 3 × 3 mm).

### Preprocessing

Preprocessing was performed using scripts generated by AFNI’s afni_proc.py and other AFNI functions (https://afni.nimh.nih.gov/) [64,65]. The first five volumes of each functional run were discarded. Subsequently, data from each scanning session were initially registered to the last volume of the final run within that session. The same T1-weighted image was used for both sessions (T1 from session 2 for 20 participants, and T1 from session 1 for the remaining participants). The aligned functional data of participants were then registered to their corresponding T1 images, with manual verification of alignment for each participant. Finally, the registered data were motion corrected and detrended (linear, quadratic and cubic) using the 3dDeconvolve function in AFNI.

### ROI definition

In the ROI-based analysis, we mainly focused on three extensively studied regions of interest (ROIs) associated with WM: early visual cortex (EVC), intraparietal sulcus (IPS) in the posterior parietal cortex, and superior precentral sulcus (sPCS) in the lateral frontal cortex. These anatomical ROIs were identified by a probabilistic atlas [66]. EVC (comprising bilateral V1, V2, and V3), IPS (comprising bilateral IPS0-5), and sPCS (comprising bilateral FEF) masks were generated by warping masks from the probabilistic atlas to each participant’s anatomical image in native space. Subsequently, the functional ROIs were identified using general linear models (GLMs), incorporating six head-motion-related regressors and three task-epoch-related regressors. The task-epoch-related regressors, consisting of sample (4.5 s), delay (9 s), and response (2 s), were modeled using boxcar functions convolved with a canonical hemodynamic response function (HRF). Functional ROIs were first determined separately for each task. For EVC, the 700 most active voxels during the sample period were selected; for IPS and sPCS, the 700 most active voxels during the delay period were used. Final functional ROIs for analysis were defined as the intersection of the top 700 voxels across both tasks within each anatomical ROI (see average ROI sizes in Table S1). In addition to the sPCS, we also used the HCP atlas to parcellate subregions across the broader frontal cortex [36,37], as detailed in Table S1.

### Representational similarity analysis

We used cross-validated representational similarity analysis (RSA) [67] to decode stimulus information within each task as well as to evaluate generalization of stimulus representation across tasks. Within-task stimulus decoding was conducted for each condition and each task separately. For a specific condition, the data were split into four folds, with a balanced number of trials for each stimulus (except for the participant who did not complete all runs). Within each fold, we averaged neural data across all trials for each stimulus to obtain pseudo-trials of stimulus-specific activity patterns [68,69]. Noise normalization was then applied to ROI analyses in Figure 2 and corresponding supplemental figures to improve reliability [70,71]. To construct the neural representational dissimilarity matrix (RDM), we computed the Pearson correlation distance between each pair of stimuli by comparing the averaged data from 3 folds with the data from the remaining 4th fold to obtain a 6×6 neural RDM. The cross-validation procedure allowed us to calculate the correlation distances along the diagonal of the neural RDM, which represented the dissimilarity between trials of the same stimulus. The lower and upper triangles of the neural RDM were averaged. Finally, we computed the Spearman correlation between the neural RDM and a continuous model RDM (which assumed that neural dissimilarity increased as a function of stimulus similarity; see Figure 2B-C upper panel). This procedure was repeated 100 times, and decoding performance was obtained by averaging the correlation coefficients across all 100 repetitions.

To investigate whether neural representations could be generalized across tasks in Experiment 1, we applied a similar analytical approach as described above. Specifically, we calculated the neural RDM by comparing location data with object data, then computed the Spearman correlation between the neural RDM and the model RDM. To maintain consistency with within-task decoding, we calculated two neural RDMs: one using 3 folds of location data and 1 fold from object data, and the other using 3 folds from object data and 1 fold from location data. The average of these two correlations was used as the measure of generalization across tasks.

### SVM decoding

We used support vector machine (SVM) decoders to examine both rule and stimulus representations for both within-task and cross-task comparisons. Decoding followed the RSA procedure, except that the neural similarity measures were replaced with SVM decoding. Specifically, we trained a classifier on 3-fold data and tested it on 4th fold, using binary C-support vector classification (C-SVC) with a linear kernel from the LIBSVM toolbox [72].

For rule decoding, we focused on the manipulation trials and excluded trials with the largest cue magnitude (cue ±180° and step 3) to avoid potentially inconsistent strategies. Moreover, we decoded three different dimensions of rule information: direction (60° & 120° vs. -60° & -120°), magnitude (±60° vs. ±120°), and identity (60° vs. -60° vs. 120° vs. -120°).

### Searchlight analysis

To examine stimuli and rule representations at the whole-brain level, we performed RSA and SVM decoding using a searchlight procedure [73]. We first warped each participant’s data in the original space to the MNI template and applied a gray matter mask [74]. The searchlight analysis was performed on the average BOLD activity across the delay period (9 – 12 s after trial onset; TR 7-9) using a spherical searchlight (radius = 9 mm), implemented using the sphere_searchlight class in the PyMVPA toolbox [75]. Results from the whole-brain searchlights were displayed on the cortical surface reconstructed with FreeSurfer [76,77] and visualized with SUMA in AFNI.

### Characterization of generalization between tasks

We used two different approaches to characterize the level of generalization between tasks [78]. The first approach requires a fixed one-to-one mapping between variables of different tasks. Similar neural representations between corresponding variables between tasks, typically quantified by means of cross-decoding and such, serve as evidence for successful generalization between these variables. In our study, this approach was implemented using cross-task RSA or SVM in Experiment 1.

When fixed mapping is not available, such as in Experiment 2, alternative approaches based on state space analysis can be employed, such as subspace overlap analysis [38] and subspace separation analysis [39,40]. These methods characterize generalization by identifying neural correlation patterns between tasks within a common subspace. All subspace analyses below focused on averaged data from the delay period.

### Subspace overlap analysis

For subspace overlap analysis, we adapted methods from previous literature [38] for fMRI data. To obtain the object subspace for stimulus information, for a specific condition, the object data were averaged across all trials for each of the six stimuli, to create a data matrix (6 stimuli × N_voxels_, where N_voxels_ refers to the number of voxels). After z-scoring the data along the stimulus dimension, we used the eig function in MATLAB to perform eigen decomposition on the covariance matrix (N_voxels_ × N_voxels_) of the data, yielding the eigenvalue matrix *V* and eigenvector matrix *W* (N_voxels_ × N_voxels_).

*V* is a diagonal matrix where the diagonal elements represent the eigenvalues corresponding to each principal component (PC). Each column in *W* represents the weights of each PC, sorted by its explained variance in a descending order. We selected the top n dimensions (n = 5 in Figure 4B-D; n = 3 in Supplementary Figure 11A-C) as the object subspace, and calculated explained variance of object data in object subspace (*Obj_ObjPC_*). The same procedure was performed to obtain the location subspace and calculated explained variance of location data (*Loc_LocPC_*). Next, we projected the averaged location data into the object subspace and calculated the variance explained by object subspace (*Loc_ObjPC_*), and projected object data into the location subspace to obtain *Obj_LocPC_*. The alignment index was defined as:

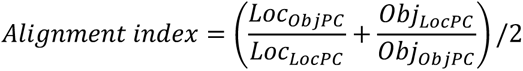

Calculation of the alignment index for cue direction followed a similar procedure. Because there were only two cue directions, trials for each cue direction were averaged into four pseudo-trials instead of one, creating eight pseudo-trials in total for the analysis.

To assess whether the subspace faithfully represented relevant information in the projected data, we projected the non-averaged (single-trial) data of one task into the subspace of the other task, and performed RSA analyses on the projected data. The RSA analyses followed previous procedures except that data were z-scored before calculating the neural RDM. Results from cross-projections were averaged across tasks.

### Subspace separation analysis

The above subspace overlap analysis characterizes neural overlap between tasks, without specifying the nature of this overlap. Theoretically, if two tasks do not overlap at all, data from the two tasks should occupy unique subspaces for each task that allow for task-specific representations. However, successful generalization between tasks should result in a shared subspace that represents both location and object tasks. We adapted the subspace separation analysis from previous literature [40] to examine whether a shared subspace existed that could represent both tasks. This analysis aimed to identify three orthogonal subspaces: one shared by both tasks and two that are unique to each individual task. Detailed steps for identifying these subspaces are described below.

The first step involved identifying a latent space that jointly represented the location and object data. Data from location (*Loc*) and object (*Obj*) tasks were split into four folds following the RSA procedure, resulting in 24 trials per task. Two data matrices (*Loc_raw_* and *Obj_raw_*), both with size of 72 × N_voxels_ (72 = 24 trials × 3 delay TRs), were constructed. To obtain latent space, PCA was performed on *Loc_raw_* and *Obj_raw_* separately, and *Loc_PreLt_* and *Obj_PreLt_* were obtained by retaining components that explained 99% variance for location and object task, respectively. Singular value decomposition was then performed on [*Loc_PreLt_*, *Obj_PreLt_*] (brackets indicate concatenation of two matrices). The left singular vectors define the latent space (N_voxels_ × N_latent_) that represents both tasks. To project task data into this latent space, *Loc_raw_* and *Obj_raw_* were averaged across TRs (24 trials × N_voxels_), and data were z-scored along the trial dimension for each task independently, resulting in *Loc_zsc_* and *Obj_zsc_*, respectively. Next, *Loc_zsc_* and *Obj_zsc_* were projected to latent space, yielding *Loc_latent_*, *Obj_latent_* (24 trials × N_latent_), and *LocObj_latent_* (48 trials × N_latent_).

The second step was to obtain unique subspaces of location and object tasks. Eigen decomposition was performed on the covariance matrix (N_latent_ × N_latent_) of *Loc_latent_*, yielding the eigenvalue matrix *V_loc_* and eigenvector matrix *W_loc_*. The first N_loc_ dimensions that captured over 99% of the explained variance were selected as the location meaningful subspace, with the remaining dimensions assigned as the location null subspace (*Loc_null_*: N_latent_ × N_loc-null_). Varying the threshold (from 99% to 95%) yielded similar results (Supplementary Figure 11D). Next, *Obj_latent_* was projected into *Loc_null_* to obtain *Obj_loc-null_* (24 trials × N_loc-null_), and eigen decomposition was performed on the covariance matrix (N_loc-null_ × N_loc-null_) of *Obj_loc-null_*, yielding the eigenvalue matrix *V_obj, loc-null_* and eigenvector matrix *W_obj, loc-null_* (N_loc-null_ × N_loc-null_). *ObjPC_loc-null_* (N_loc-null_ × N_obj-unique_) was then obtained by discarding the trailing dimensions that capture less than 1% variance of *Obj_latent_*. *Loc_null_* and *ObjPC_loc-null_* were multiplied to produce a single orthonormal subspace *Z_obj-unique_* (N_latent_ × N_obj-unique_), which captured object-specific variance while excluding location-related variance. *LocObj_latent_* was projected into *Z_obj-unique_* to obtain the object-unique responses, *Y_obj-unique_* (48 trials × N_obj-unique_). The same procedure was performed to obtain location-unique responses *Y_loc-unique_* (48 trials × N_loc-unique_). Finally, two orthonormal unique subspaces, *Q_loc-unique_* (N_latent_ × N_loc-unique_) and *Q_obj-unique_* (N_latent_ × N_obj-unique_) were obtained through gradient descent manifold optimization (Stiefel manifold) using the Manopt toolbox [79]. The goal of this optimization was to minimize the sum of squared residuals between [*Y_loc-unique_*, *Y_obj-unique_*] and [*LocObj_latent_* · *Q_loc-unique_*, *LocObj_latent_* · *Q_obj-unique_*], subject to Q_loc-unique_ ⊥ Q_obj-unique_.

The third step was to obtain the shared subspace. The primary shared subspace *Qpri_shared_* (N_latent_ × [N_latent_ - N_loc-unique_ - N_obj-unique_]) was obtained by computing the null space of [*Q_loc-unique_*, *Q_obj-unique_*]. *LocObj_latent_* was then projected into *Qpri_shared_*, and PCA was applied to obtain *Qmini_shared_* ([N_voxels_ - N_loc-unique_ - N_obj-unique_] × N_shared_). Finally, the shared subspace *Q_shared_* (N_latent_ × N_shared_) was obtained by multiplying *Qpri_shared_* and *Qmini_shared_*. The shared subspace *Q_shared_* contained dimensions that were either meaningful for both tasks or non-meaningful for either task. The non-meaningful dimensions, by definition, do not contribute to either task and thus do not affect the assessment of shared information.

Lastly, *Loc_latent_* and *Obj_latent_* were projected into unique and shared subspaces, decoding and multi-dimensional scaling (MDS) analysis were performed within each subspace. MDS analysis based on Euclidean distances was conducted using the cmdscale function in MATLAB. This subspace separation procedure was repeated 100 times and averaged across 100 repetitions for the final results.

### Statistical analyses

To quantify the linear trend between cue magnitude and behavioral performance, we performed a linear mixed-effects model (LMEM) analysis on cue magnitude (excluding 180° in the location task and step-3 in the object task) and behavioral performance (accuracy or reaction time). We used the fitlme function in MATLAB:

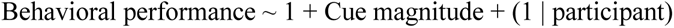

To evaluate strategy changes in the object task between Experiment 1 and 2, we modeled behavioral performance across all cue magnitudes:

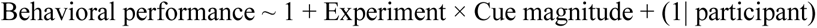

For all LMEMs, we evaluated *β* coefficients and p-values of the task variables, and used the anova function to obtain the F-values of main and interaction effects of experiment. Paired t-tests were used to evaluate differences between cues, and the resulting two-tailed p-values were corrected for multiple comparisons using False Discovery Rate (FDR).

To assess statistical significance of neural decoding results, we used a sign-flip permutation procedure. For a specific decoding result, the p-value was computed by comparing the true decoding result against a null distribution. The null distribution was generated by randomly multiplying either 1 or -1 to individual decoding results and averaging the sign-flipped results across all participants for 10000 times, resulting in a null distribution of 10000 decoding results. The resulting one-tailed p-values were corrected for multiple comparisons using FDR across conditions or ROIs, unless specified. A similar procedure was applied to assess the statistical difference between conditions within the same experiment and between experiments (e.g., alignment index). The null distribution for between-experiment comparisons was generated by randomly splitting results of all participants from two experiments (n = 46) into two groups and calculating the difference between groups for 10000 times.

To examine the significance of searchlight analyses, we used the 3dttest++ function in AFNI to perform t-tests for each condition and condition differences. To reduce computational load, we used a two-stage procedure for cluster-based permutation [80]: First, we sign-flipped the whole-brain results of each participant for 100 times to obtain individually randomized results. Second, the randomized results were bootstrapped from each participant and subjected to one-tailed t-tests (α = 0.05) across participants to obtain a t-value map. The second step was repeated 10000 times, and the sum of t-values in the largest cluster from each bootstrap was taken to create a null distribution. The contiguous t-value clusters of true results and the null distribution underwent multiple comparisons correction, and the obtained p-values were corrected using Family-Wise Error Rate (FWER) method with a threshold of α = 0.01.

## Acknowledgements

This work was supported by the Strategic Priority Research Program of the Chinese Academy of Sciences (XDB1010202), the Ministry of Science and Technology of China (STI2030-Major Projects 2021ZD0203701, 2021ZD0204202), the National Natural Science Foundation of China (32271089), and CAS Project for Young Scientists in Basic Research (YSBR-071) to Q.Y..

## Supplemental Figures

**Figure S1.**
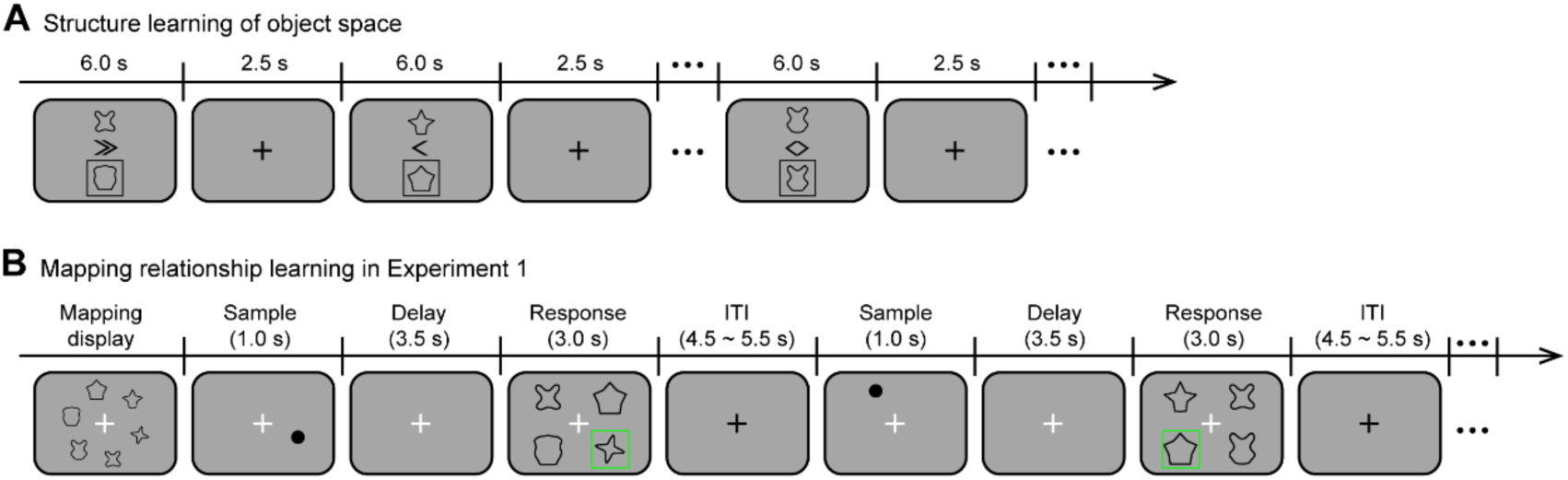
Schematics for object-space learning and mapping learning. (A) Structure learning of object space in Experiments 1 and 2. During the behavior training of object task, participants acquired the structure of object space by learning the transitional relations between objects drawn from the space. Specifically, on each trial, participants saw a pair of two objects presented on the screen for 6 s, one above fixation and one below fixation. A cue was presented at the center, indicating that manipulating the object above according to the cue would result in the object below (outlined by a square). There were 21 trials in each run, with object pairs randomly chosen from all possible transitional relations in the object space. Participants were instructed to continue learning the translational relations until they could draw the correct relationships between objects. Participants on average performed 1 - 3 runs for the structure learning task. (B) Learning of mapping between object and location spaces in Experiment 1. In Experiment 1, participants additionally learned the mapping between the two stimulus spaces prior to scanning. At the beginning of each run, the mapping relationships between objects and locations were shown on the screen (each object presented in its corresponding location). Participants pressed the space key to move onto the test stage if they believed they had remembered the mapping relationships. On each test trial, a black dot appeared on the screen for 1 s, followed by a delay of 3.5 s. During the response period, participants needed to recall the object corresponding to the dot location and choose the correct answer from four options within 3 s. There were 30 trials in each run. The mapping learning ended when participants’ performance reached > 90%.

**Figure S2.**
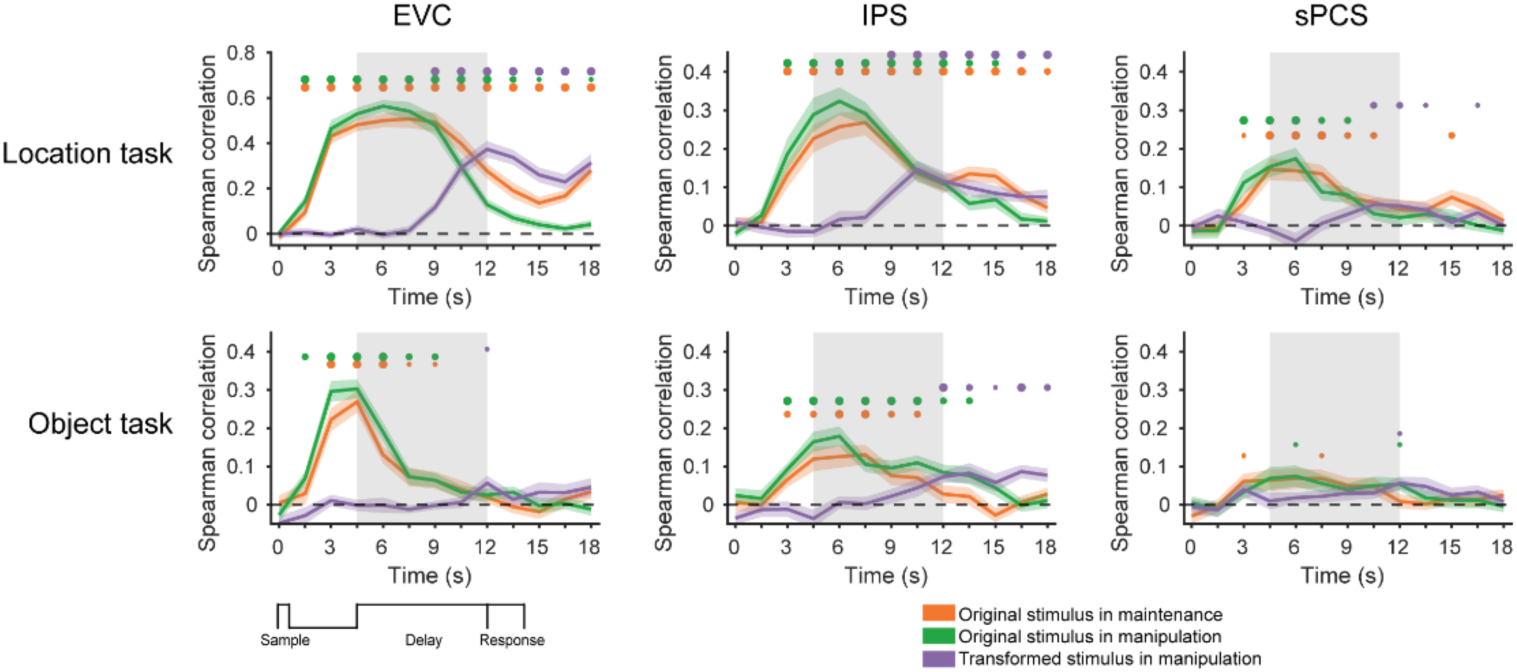
Control analysis for the anticorrelation effect in within-task RSA in Experiment 1. To remove the anticorrelation between the original and transformed stimuli in the manipulation condition, we added 20% of maintenance trials to the manipulation data, ensuring equaled trial numbers for all original-transformed stimulus combinations. We also added these maintenance trials to the maintenance data to maintain equaled trial numbers across conditions. During the trial-balancing process, we ensured that the same trials were not reused in training and testing. After trial-balancing, the early decoding of the transformed location in EVC disappeared.

**Figure S3.**
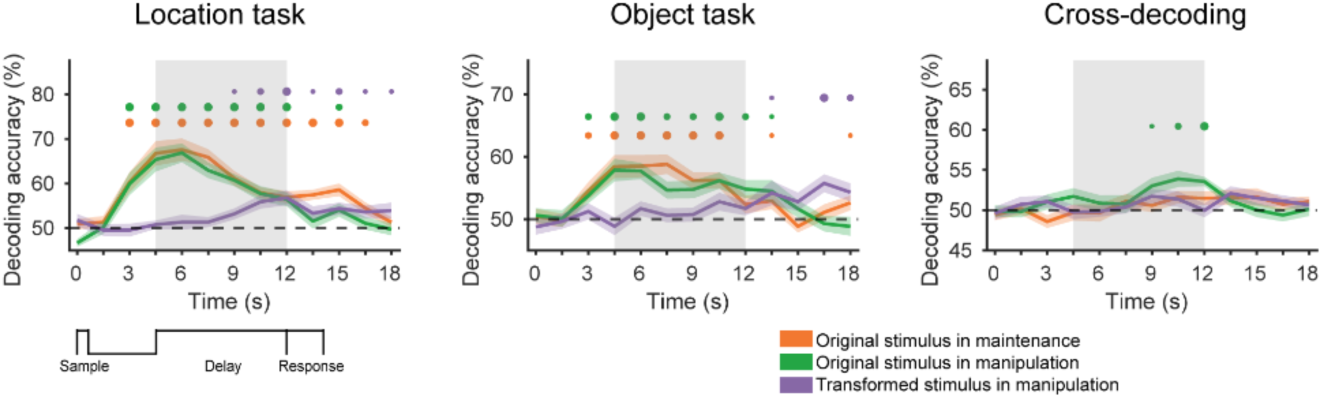
Within- and cross-task SVM decoding results in the IPS for Experiment 1. The within- and cross-task SVM decoding results in the IPS remained qualitatively similar to those using RSAs, suggesting that the observed results in the IPS were not specific to the method being used.

**Figure S4.**
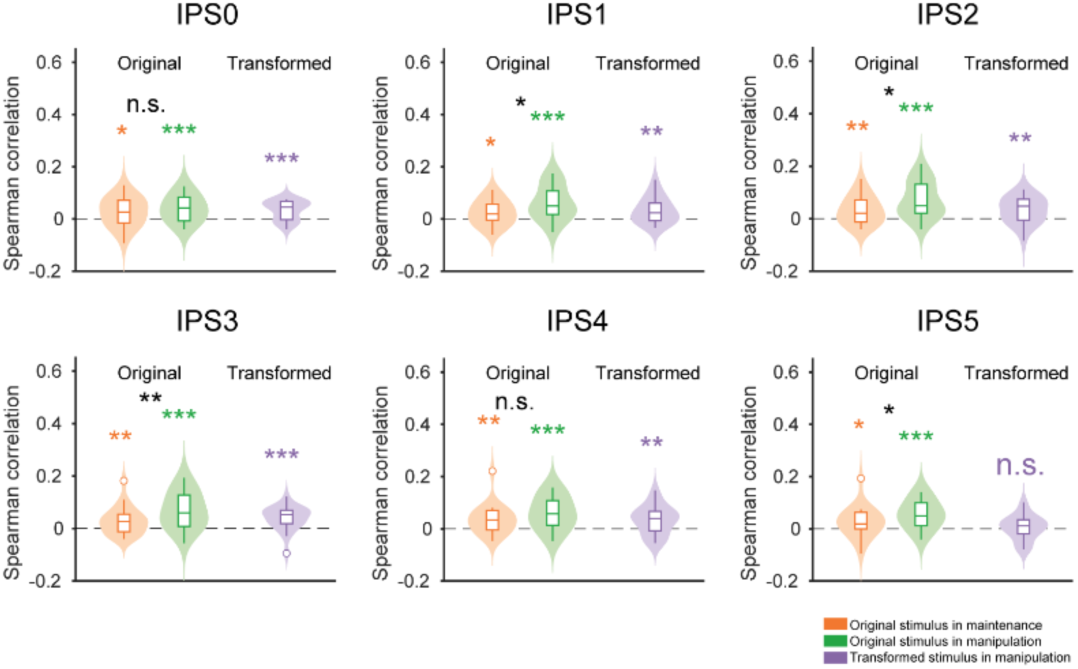
Cross-task RSA results in IPS subregions for Experiment 1. The cross-task RSA results largely remained in different subregions of the IPS, suggesting that the observed results were robustly present in the IPS.

**Figure S5.**
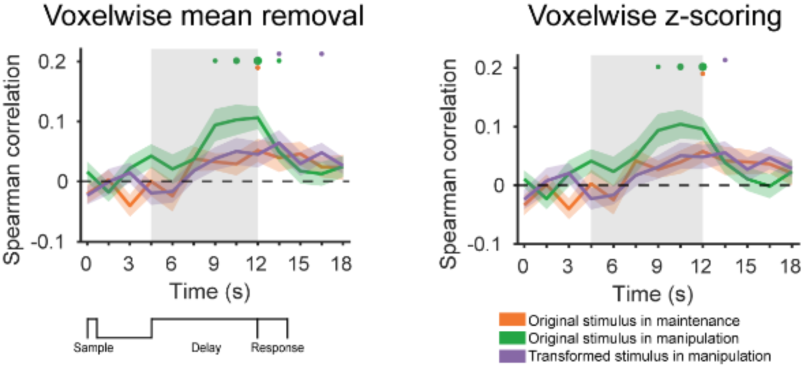
Cross-task RSA results in the IPS after removing activation differences between conditions in Experiment 1. Left panel shows results after removing mean activity of each condition at the voxelwise level, and right panel shows results with voxelwise z-scoring within condition. The cross-task RSA results remained largely unchanged, suggesting that the enhanced decoding performance for the manipulation condition in the IPS cannot be explained by activation differences between conditions.

**Figure S6.**
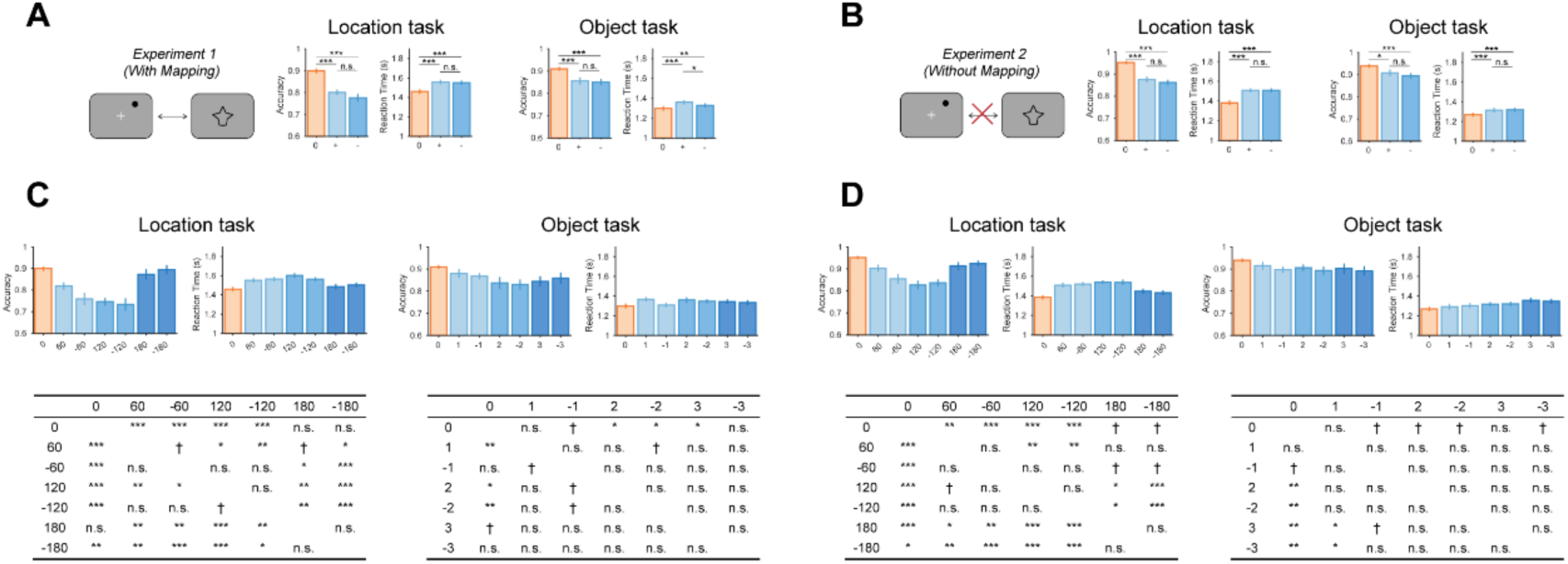
Behavioral performance plotted as a function of cue direction and cue identity. (A) Behavioral accuracy and reaction time in the location (middle panels) and object tasks (right panels) under different cue directions in Experiment 1. ‘+’ and ‘-’ indicate clockwise and counterclockwise directions, respectively. (B) Same as (A) but with results from Experiment 2. (C) Behavioral accuracy and reaction time in location and object tasks under different cue identities in Experiment 1. The tables below show the p-values of pairwise comparisons. Upper triangles indicate p-values for behavioral accuracy, and lower triangles indicate p-values for reaction time. (D) Same as (C) but with results from Experiment 2. Black asterisks indicate FDR-corrected significance, \*\*\**p* < 0.001, \*\**p* < 0.01, \**p* < 0.05, †*p* < 0.1, n.s. *p* ≥ 0.1. Error bars denote ±1 SEM.

**Figure S7.**
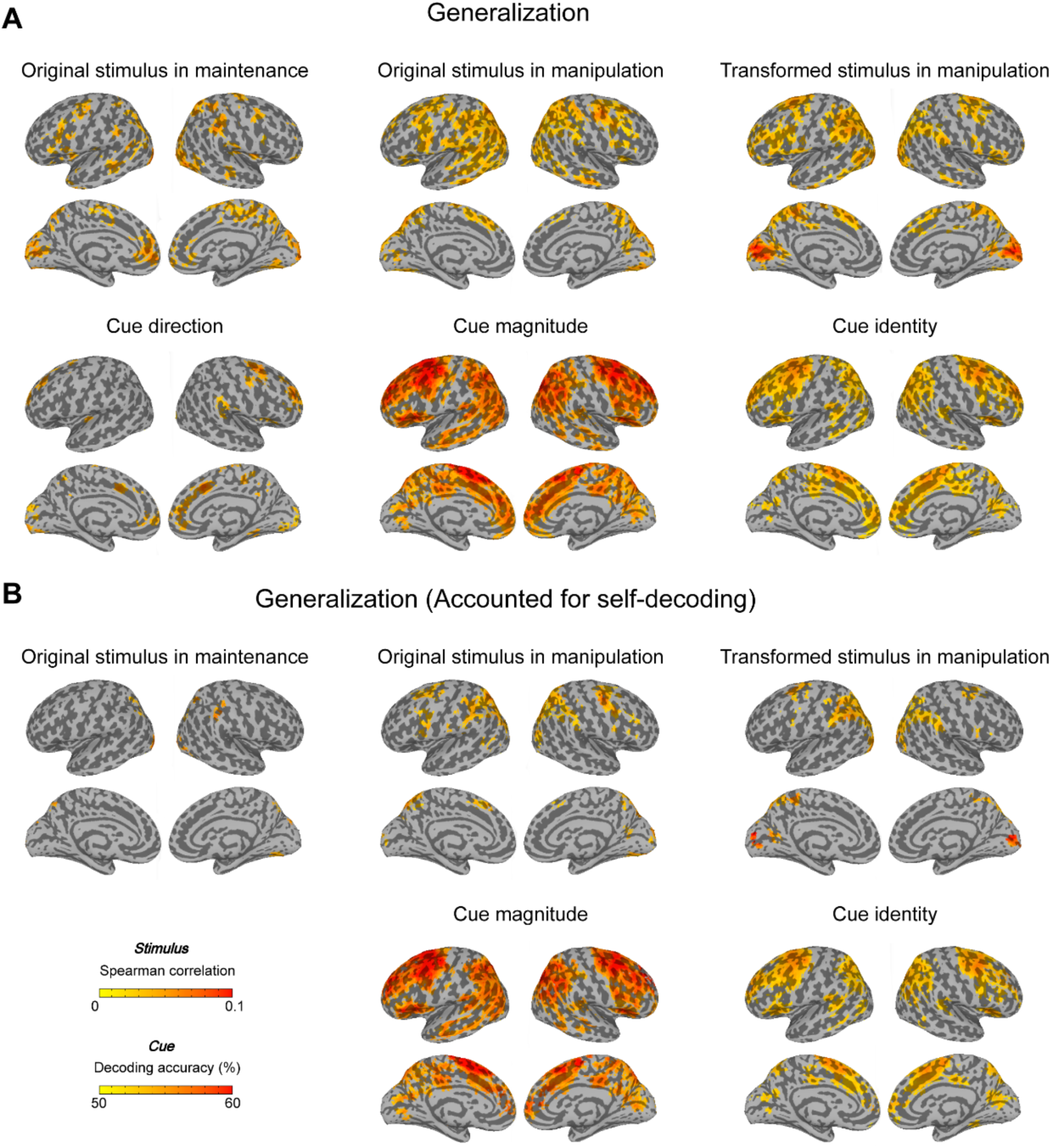
Cross-task decoding searchlight results in Experiment 1. (A) Cross-task searchlight results for each stimulus type (first row) and for different rule dimensions (second row). (B) Same as (A), but results were thresholded using self-decoding results, such that only regions with significant within-task decoding were displayed. After thresholding, the searchlight results were largely consistent with ROI analyses, showing that stimulus generalization during WM manipulation primarily engaged the IPS and frontal subregions such as the sPCS. Nevertheless, it is important to consider the signal-to-noise ratio (SNR) of fMRI signals in different brain regions when interpreting these findings, because the generalizability of signals in a given brain region depends at least partly on the strength of within-task stimulus representations. It is possible that some higher-order or deep brain regions also represent WM information, but their signals are too weak to be detected as significant. Relatedly, note that although subregions in the medial network, including medial PFC and ACC, demonstrated higher stimulus generalization for WM maintenance compared to manipulation, within-task stimulus decoding in these regions was not significant. One explanation is that WM representations in the medial network were generally weak and susceptible to low SNR, but because noise in the two tasks may have differed in direction, cross-decoding could be less affected by within-task noise. This hypothesis requires further investigation, but if true, it would indicate that our generalization approach may help mitigate the influence of low SNR at the resolution of fMRI.

**Figure S8.**
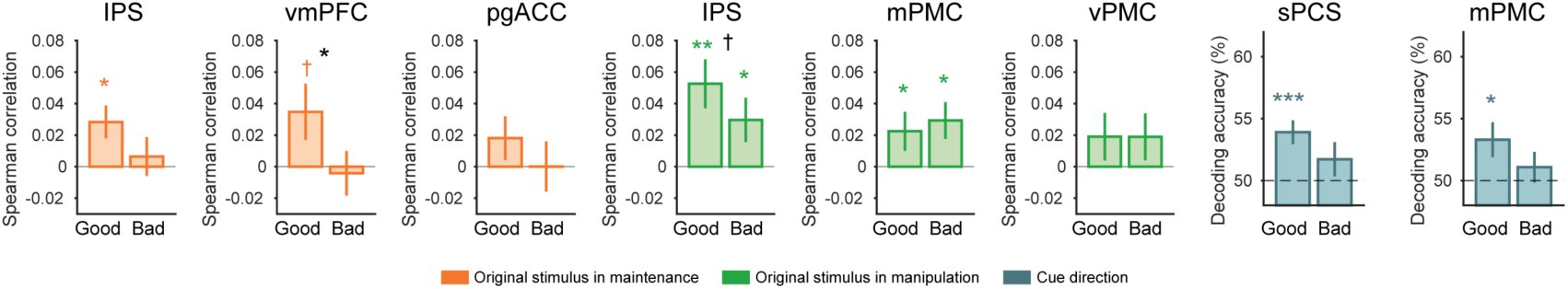
Neural correlates of behavioral performance in cross-task decoding for Experiment 1. Cross-task decoding results for stimuli and rules in IPS and frontal subregions, compared between trials with fast (good) and slow (bad) reaction times (RTs). Only ROIs and conditions with significant original stimulus or rule decoding are shown. fMRI data were median-split based on reaction times, and cross-decoding was performed separately for each half of the data. Asterisks indicate FDR-corrected significance, \*\*\**p* < 0.001, \*\**p* < 0.01, \**p* < 0.05, †*p* < 0.1, n.s. *p* ≥ 0.1. Error bars denote ±1 SEM.

**Figure S9.**
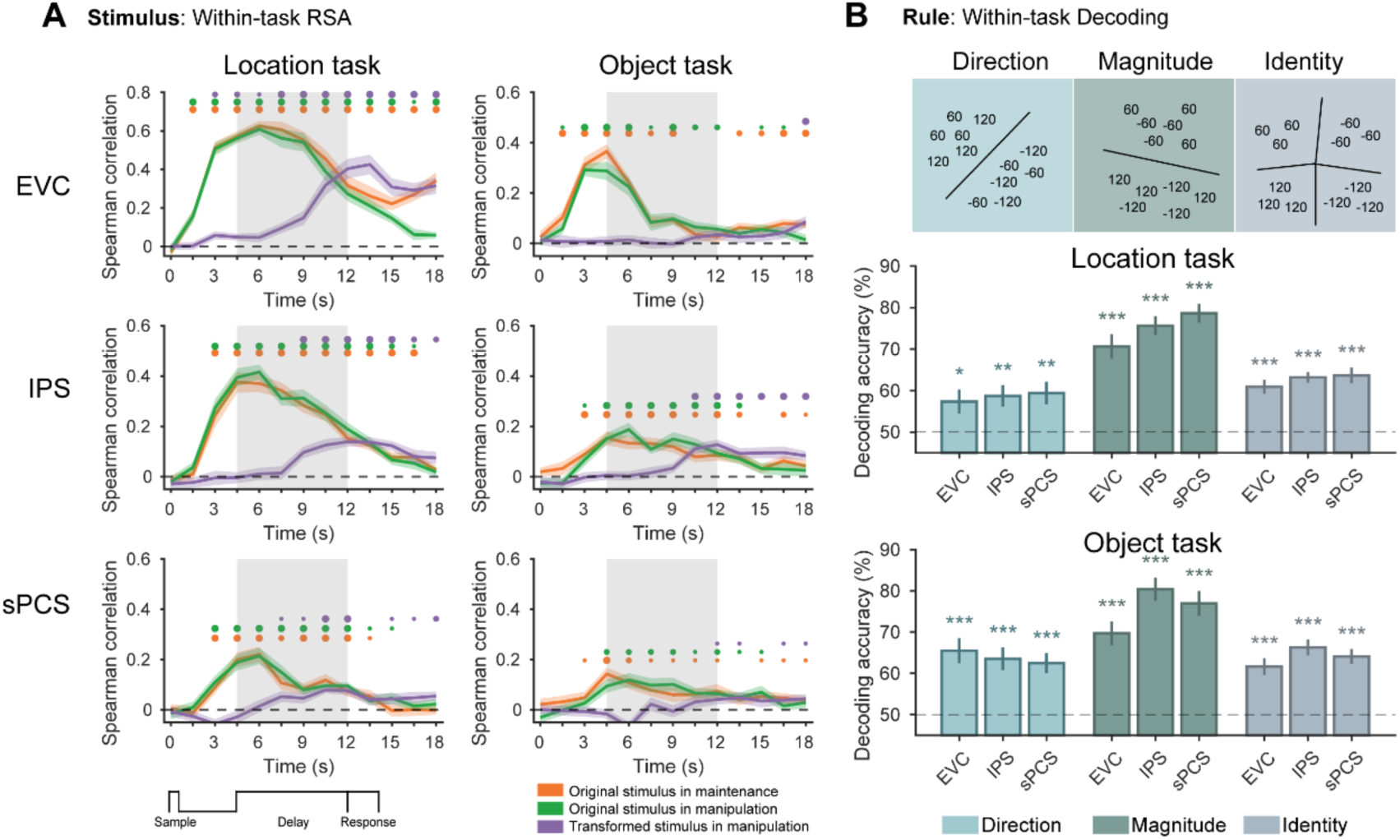
Within-task decoding in Experiment 2. (A) Within-task RSA results for different stimulus types. (B) Within-task SVM decoding results for different rule dimensions.

**Figure S10.**
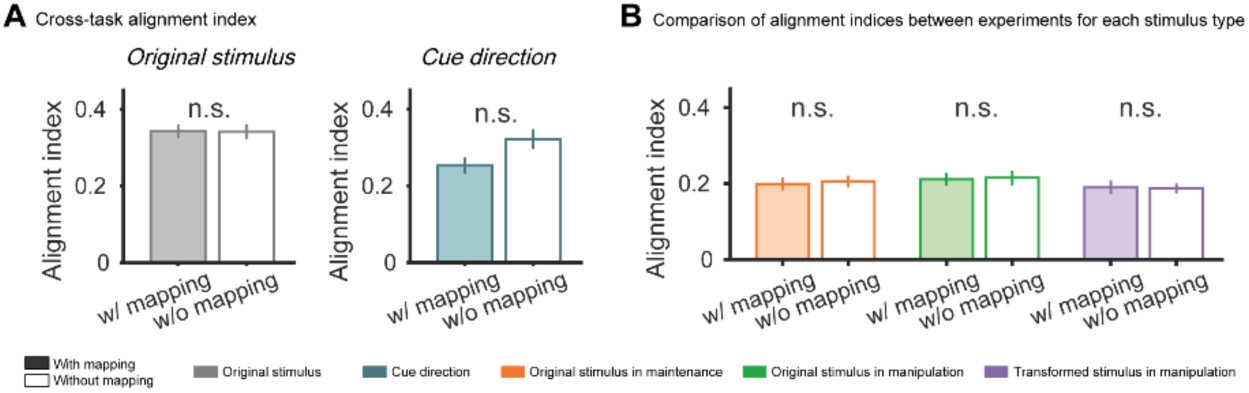
Alignment index from subspace overlap analysis in the EVC. No significant difference was observed between experiments in EVC. w/ mapping: with mapping (Experiment 1); w/o mapping: without mapping (Experiment 2).

**Figure S11.**
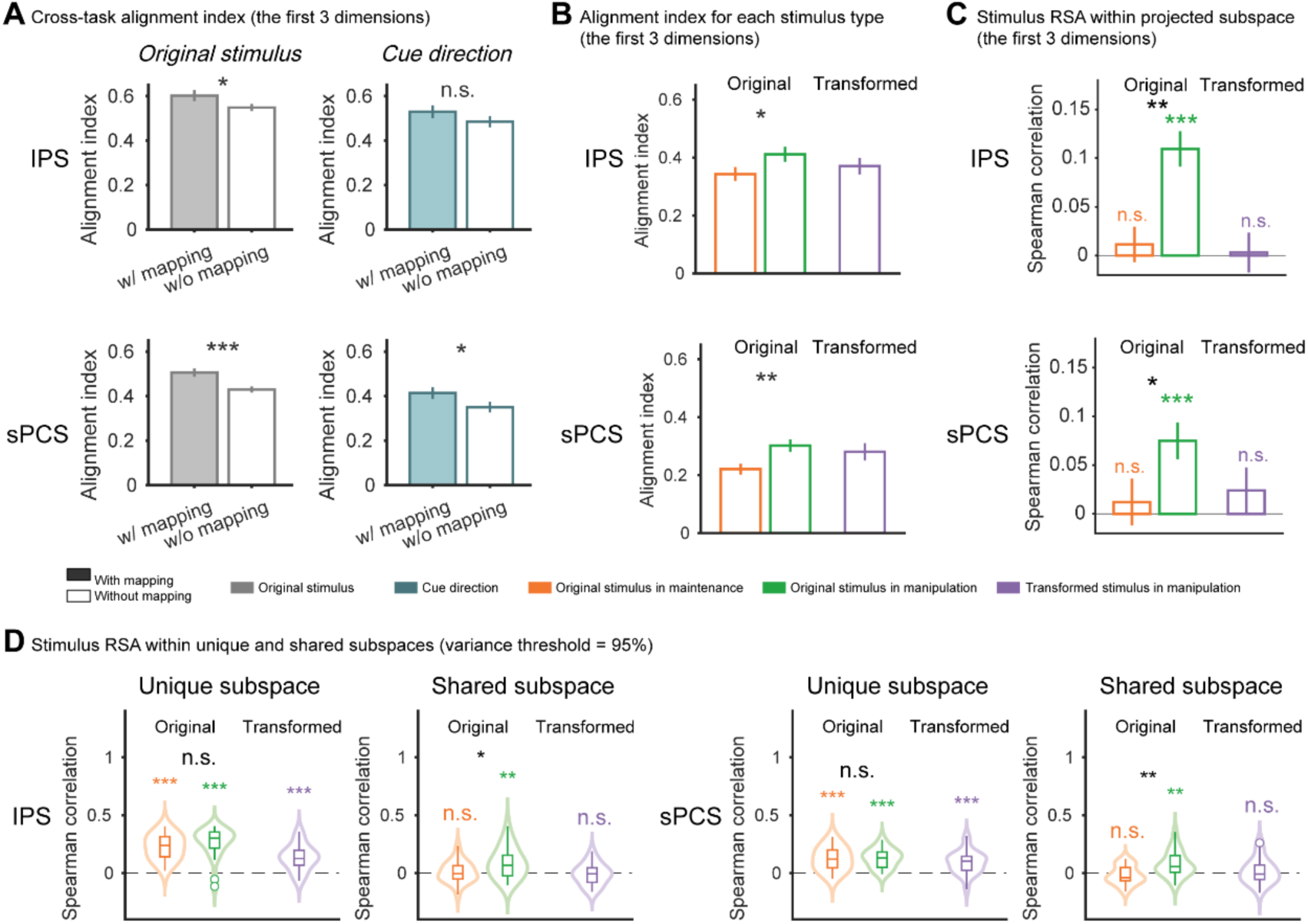
Control analysis for subspace overlap and subspace separation analyses. (A) Alignment indices for the first 3 dimensions between location and object tasks in Experiment 1 (with mapping) and 2 (without mapping) in IPS and sPCS. Left panel shows results combined across the original stimulus conditions, while right panel shows results for cue direction. (B) Alignment indices for each stimulus type using the first 3 dimensions in Experiment 2. (C) Within-task stimulus RSA results of subspace overlap analysis using first 3 dimensions in Experiment 2. Results remained qualitatively similar to those obtained with the first 5 dimensions. (D) Within-task stimulus RSA results (average across location and object tasks) in the unique and share subspaces in IPS and sPCS from Experiment 2. The variance threshold used to define the subspaces was 95%. Results were consistent with the main findings.

**Figure S12.**
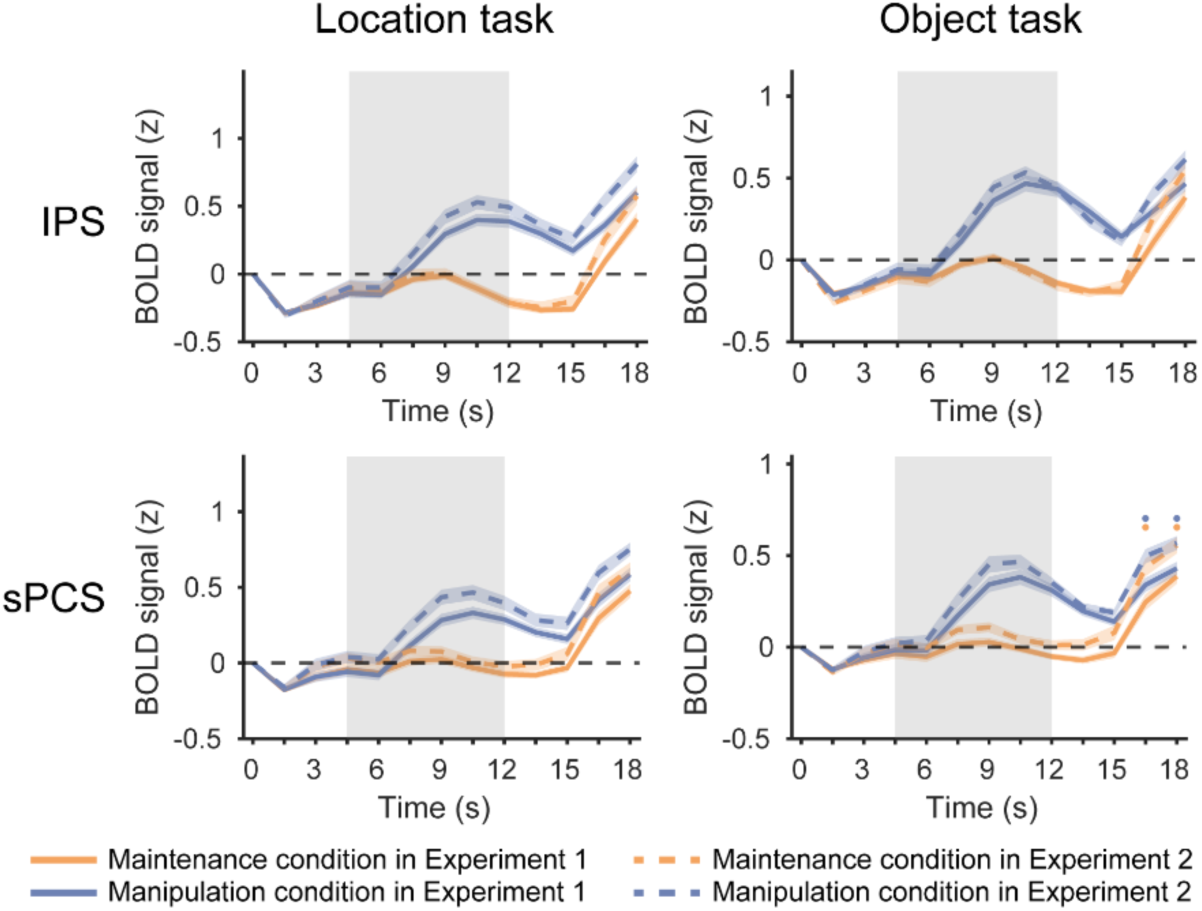
Time course of BOLD signal changes in the IPS and sPCS. Time course of BOLD signal changes in the IPS and sPCS. Horizontal dots on top indicate FDR-corrected significance for the difference between experiments over time (orange dots: difference in maintenance conditions; blue dots: difference in manipulation conditions), with dot sizes indicating different levels of significance: large (*p* < 0.001), medium (*p* < 0.01), and small (*p* < 0.05). Error bars denote ±1 SEM. Although manipulation conditions evoked larger BOLD signal changes compared with maintenance conditions, BOLD activity between the two experiments remained largely comparable during the delay, suggesting that differences in the alignment index cannot be explained by differences in BOLD activity.

**Figure S13.**
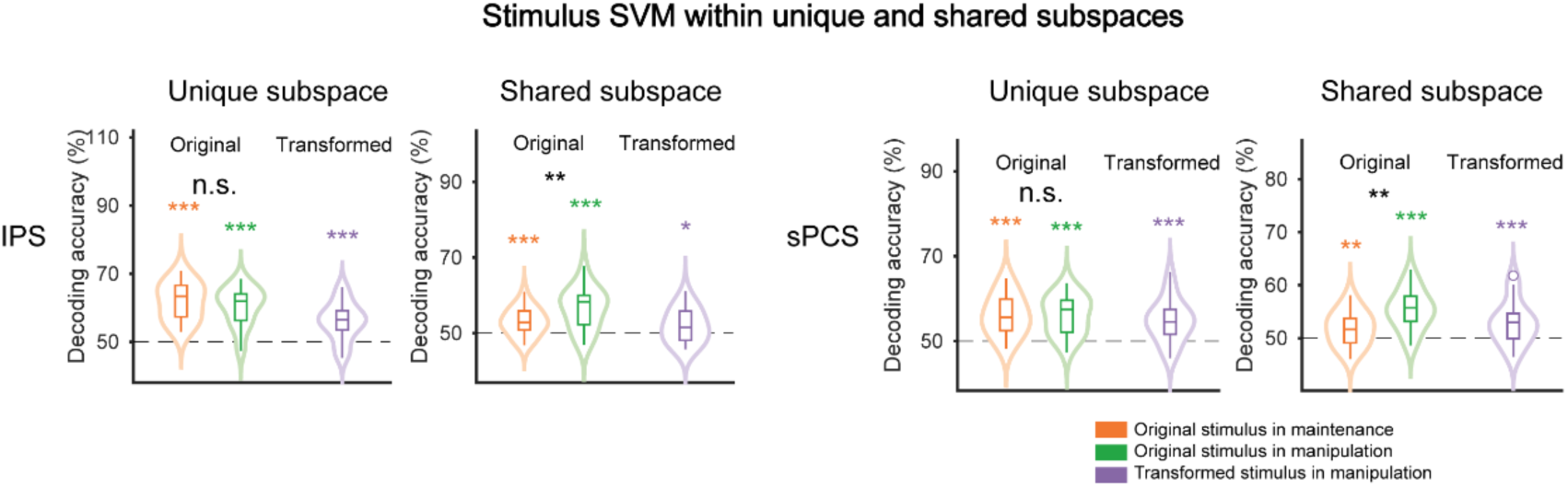
Stimulus SVM results within unique and share subspaces in IPS and sPCS of Experiment 2. SVM Results in the subspace separation analyses remained qualitatively similar to those using RSA.

**Figure S14.**
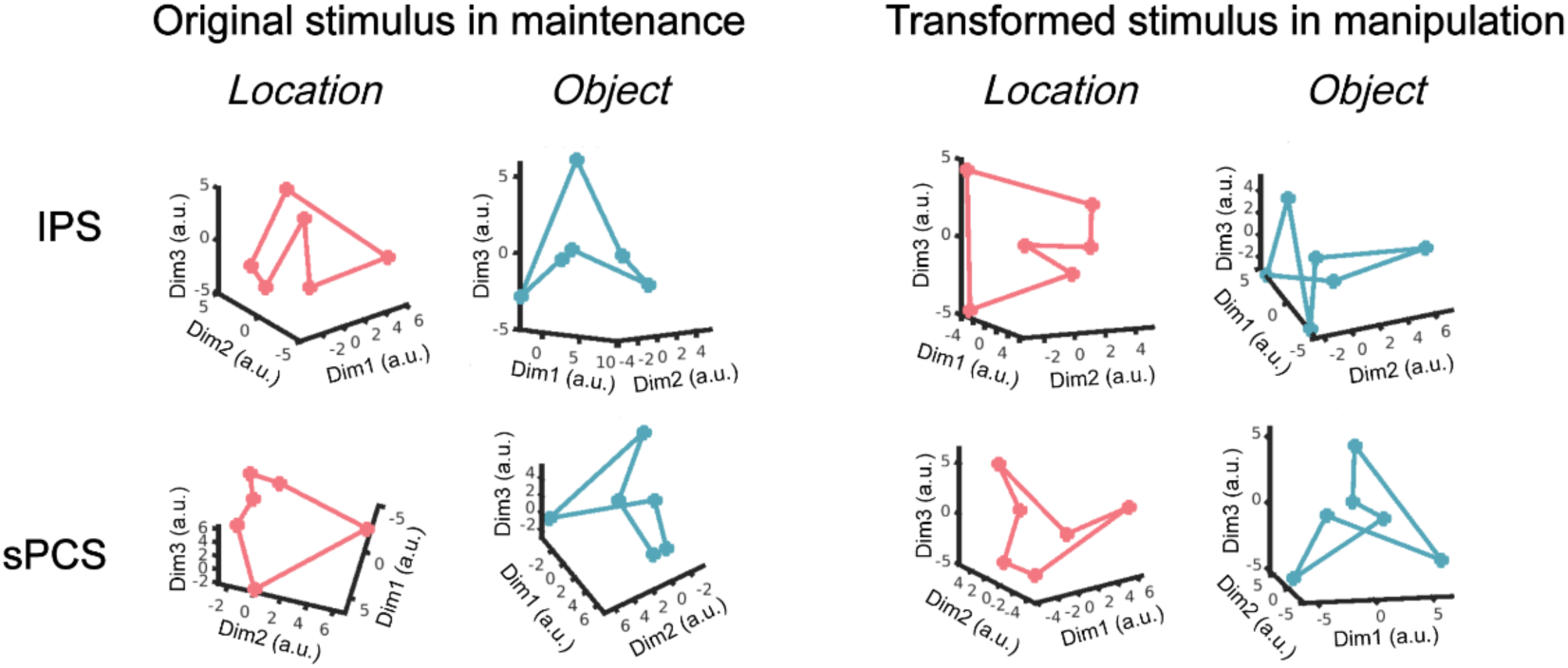
MDS visualization of the original stimulus in maintenance and transformed stimulus in manipulation in IPS and sPCS for Experiment 2.

**Figure S15.**
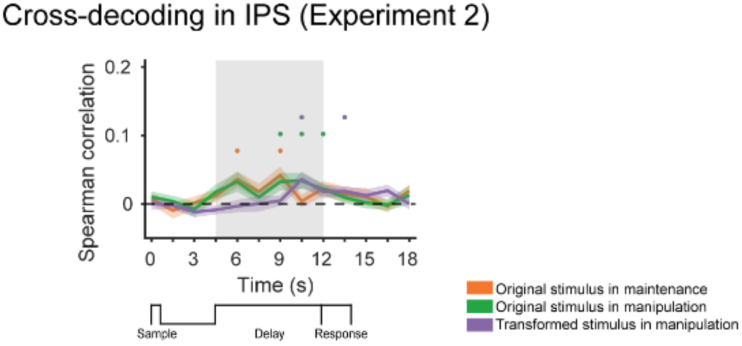
Cross-task RSA results in IPS for Experiment 2. Since there was no explicit one-to-one mapping between location and object stimuli in Experiment 2, we used a cross-validated RSA procedure in the main analyses to identify potential mapping in the training data, which were then applied to the testing data for cross-task RSA. Specifically, there were 12 possible mapping relationships based on the task design. We calculated the average Euclidean distance of paired location and object stimuli for each mapping in the training data, and applied the one with the smallest distance to the test data.

**Table S1.**
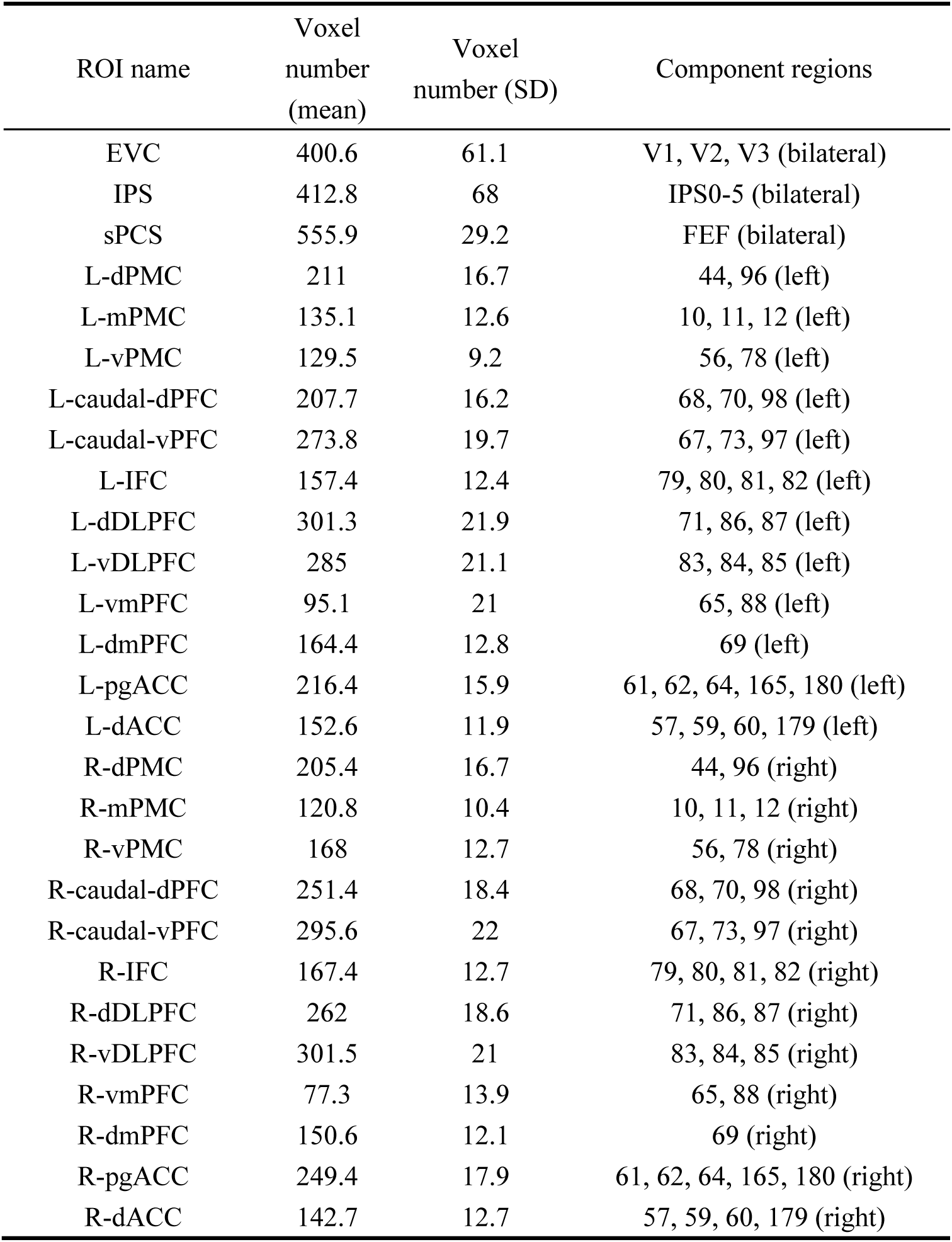
ROI definition and size in Experiment 1. Average ROI sizes across participants (mean and SD: columns 1 and 2) and component regions in the probabilistic atlas and HCP atlas for defining each ROI (column 3).

| ROI name | Voxel<br>number<br>(mean) | Voxel<br>number (SD) | Component regions |
| --- | --- | --- | --- |
| EVC | 400.6 | 61.1 | V1, V2, V3 (bilateral) |
| IPS | 412.8 | 68 | IPS0-5 (bilateral) |
| sPCS | 555.9 | 29.2 | FEF (bilateral) |
| L-dPMC | 211 | 16.7 | 44, 96 (left) |
| L-mPMC | 135.1 | 12.6 | 10, 11, 12 (left) |
| L-vPMC | 129.5 | 9.2 | 56, 78 (left) |
| L-caudal-dPFC | 207.7 | 16.2 | 68, 70, 98 (left) |
| L-caudal-vPFC | 273.8 | 19.7 | 67, 73, 97 (left) |
| L-IFC | 157.4 | 12.4 | 79, 80, 81, 82 (left) |
| L-dDLPFC | 301.3 | 21.9 | 71, 86, 87 (left) |
| L-vDLPFC | 285 | 21.1 | 83, 84, 85 (left) |
| L-vmPFC | 95.1 | 21 | 65, 88 (left) |
| L-dmPFC | 164.4 | 12.8 | 69 (left) |
| L-pgACC | 216.4 | 15.9 | 61, 62, 64, 165, 180 (left) |
| L-dACC | 152.6 | 11.9 | 57, 59, 60, 179 (left) |
| R-dPMC | 205.4 | 16.7 | 44, 96 (right) |
| R-mPMC | 120.8 | 10.4 | 10, 11, 12 (right) |
| R-vPMC | 168 | 12.7 | 56, 78 (right) |
| R-caudal-dPFC | 251.4 | 18.4 | 68, 70, 98 (right) |
| R-caudal-vPFC | 295.6 | 22 | 67, 73, 97 (right) |
| R-IFC | 167.4 | 12.7 | 79, 80, 81, 82 (right) |
| R-dDLPFC | 262 | 18.6 | 71, 86, 87 (right) |
| R-vDLPFC | 301.5 | 21 | 83, 84, 85 (right) |
| R-vmPFC | 77.3 | 13.9 | 65, 88 (right) |
| R-dmPFC | 150.6 | 12.1 | 69 (right) |
| R-pgACC | 249.4 | 17.9 | 61, 62, 64, 165, 180 (right) |
| R-dACC | 142.7 | 12.7 | 57, 59, 60, 179 (right) |

## Notes

### Competing Interest Statement

The authors have declared no competing interest.

### Summary of Updates

author affiliations, emails, and ORCID IDs updated.

